# Scaling of Protein Function Across the Tree of Life

**DOI:** 10.1101/2023.03.08.531481

**Authors:** Riddhi Gondhalekar, Christopher P. Kempes, Shawn Erin McGlynn

## Abstract

Scaling laws are a powerful way to compare genomes because they put all organisms onto a single curve and reveal non-trivial generalities as genomes change in size. The abundance of functional categories in a given genome scales with genome size, suggesting that universal constraints shape category abundance. Here we look across the tree of life to understand how genome evolution may be related to functional scaling. We revisit previous observations of functional genome scaling with an expanded taxonomy by analyzing 3726 bacterial, 220 archaeal, and 79 unicellular eukaryotic genomes. We find that for some functional classes, scaling is best described by multiple exponents, revealing previously unobserved shifts in scaling as genomes grow or contract. Furthermore, scaling varies between phyletic groups and is less universal than previously thought, with scaling shifts varying uniquely between domains and scaling exponents varying uniquely between phyla. We also find that individual phyla frequently span scaling exponents of functional classes, revealing that individual clades can move across scaling exponents. Across the tree of life, variability in functional scaling is not accounted for by genome phylogeny, suggesting that physiological and/or cell plan outweighs phylogeny.

## INTRODUCTION

Scaling relationships are ubiquitous and are suggestive that a set of rules governs how a system works (1). In biological systems, a common challenge for understanding the history of life, extant species diversity, and the possibilities for life beyond Earth is to find biological laws that systematize biology in these different contexts. As we know it on Earth today, life is unified in using a conserved set of proteins (2, 3), but organisms differ dramatically in their unique and shared protein complements (e.g. (4, 5)). Despite this diversity, a variety of scaling relationships across the tree of life have been identified, suggesting universal constraints shape processes and form across lineages, even if accomplished in evolutionarily independent ways (6–17). By identifying the connection between scaling relationships and physical constraints, it may be possible to demonstrate that they represent laws. For example, in bacteria, scaling in physiology, metabolism, and growth is often due to connections between physical constraints and cell size (9–11).

One class of scaling relationships is those observed comparing the total number of annotated proteins to the abundance of specific functional classes. Genome size is a complex trait with relationships to temperature (18), physiology (19), lineage history (20), and cell size (21). When genomes expand or contract, function may be gained or lost in a non-uniform way. Starting with a seminal study by van Nimwegen (12) and subsequently expanded by others (13, 15–17, 22–25), the number of genes in a functional category has been found to scale as a power-law with the total number of genes in the genome (12, 13).

Such scaling patterns have been demonstrated using functional annotations, which indicated that metabolic and electron transfer processes scale uniquely between clades, highlighting that different metabolisms have different regulatory and energy demands, whereas other genomically encoded processes, such as translation, scale similarly across clades (12, 13, 16). Recent work has also suggested that at the level of broad enzyme categories, there are no shifts across taxa in scaling exponents suggesting a greater universality at the level of enzyme “operations” (E.C. level 1) than observed in functional scaling (17). This sets up an interesting question about how functional categories scale, as this categorization projects the same enzymes onto physiological processes rather than broad categories of enzyme function.

With the much larger number of genomes available today, it is worth revisiting functional scaling relationships across an expanded phylogenetic diversity. As two examples of many - Thaumarchaeota genomes only became available in the 2010s (26), after functional protein scaling relationships were first reported. Metabolisms once thought to be rather restricted have now been found more broadly, for example, methane metabolism seems more widespread across the archaeal tree (see (27) and references contained therein). Across this increased genomic and physiological diversity, it may also be possible to test the effects of major evolutionary transitions in connection with known shifts in physiological scaling (10, 11) and to test trends within taxa with greater statistical power.

Here we use COG (clusters of orthologous genes) functional annotations to validate previous observations with an expanded taxonomy and find previously unobserved scaling relationships and shifts in those relationships. We make use of the eggNOG (Evolutionary genealogy of genes: Non-supervised Orthologous Groups) database (28) as a starting data set which covers more taxonomic diversity than has been used in previous studies, and we supplement this by annotating other genomes of interest using the eggNOG annotator: including members of the CPR, DPANN, Melainabacteria, Asgard archaea and some lower Eukaryotes. We find a substantial variation of functional scaling between clades and suggest that some major evolutionary transitions may have occurred in conjunction with changes in functional scaling which would mirror the physiological shifts observed over these transitions.

## MATERIALS AND METHODS

### Data Processing and Power Law Fitting

We considered COG (clusters of orthologous genes) annotations of sequenced genomes using the <u>eggNOG database</u>. Using the eggNOG-mapper v2, we annotated the proteins of groups not available in the eggNOG database, like Melainabacteria, CPR, DPANN Asgard archaea, and some unicellular eukaryotes. The genome information used for annotations using the eggNOG-mapper v2 was obtained from the NCBI server. Asgard archaeal genome information was obtained from the <u>GTDB (Genome Taxonomy Database)</u> website. We mapped the files to get their COG annotations from the eggNOG annotations. Further, we found the total number of annotations in a COG category and the total number of functional annotations in an organism. In cases where a protein had more than one COG annotation, these were split and counted as individual counts in the total number of functional annotations.

We used Ordinary Least Square (OLS) regressions of the log-transformed data. We chose this method for easy intercomparison with various previous studies that employ this method. The 95% confidence interval on slope and intercept was found using t continuous random variable of scipy.stats.linregress from SciPy.

We carried out a null test by shuffling the COG category annotations. This resulted in ortho-groups with very different abundances per organism, but the total number of COGs with categories remained the same. The power law fits for the shuffled data are in the Supplementary figures (Supplementary figure 1).

### Breakpoints

A challenge in scaling data is that often there is asymptotic behavior for the largest and smallest sizes in the system, which in some cases is anticipated from theory (9, 10, 29). We used the piecewise_regression library to compute the breakpoints. Within a taxonomic group, we present scaling analyses where either a single scaling relationship or two scaling relationships with a breakpoint are the best fits. We select the breakpoint model when it produces a significant improvement in the goodness of fit which we take to be a 5% decrease in the residual sum of squares values (RSS) compared to the single scaling relationship fit. It was calculated by dividing the RSS for one breakpoint by the RSS for no breakpoints (Supplementary Tables 7 and 8). After finding the breakpoint locations, we fit the points using the same procedure as fitting the individual data plots. To visualize the locations of the breakpoints across genomes, we overlaid the breakpoint location against the genome spans of phyla (Figure 7, Supplementary Tables 3 and 4).

### Binning

OLS fits can be dominated by ranges of x values with the most data, which can lead to inaccurate power-law fits. An uneven amount of data across the scales is definitely characteristic of our data. To deal with this unequal sampling, we binned the dataset and then performed the OLS fit to the binned data. Supplementary Figure 2 shows the results of the binned data and fits overlaid the full dataset, and Supplementary Table 5 compares the fit values where we only find negligible differences in some cases.

### Z Statistics

We calculated the Z-statistics score to quantify the significance of the deviations of individual phyla exponents from their overall domain exponents using the formula developed by (13).

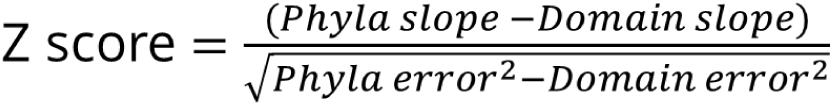

For plotting, we consider both positive and negative deviations (Figure 6 and Supplementary Table 6).

### Phylogenetic Distance Plots

We calculated the phylogenetic distances between phyla using the Dendropy library and the Hug et al., 2016 tree of life (30). Since we did a phyla-level analysis, we took the average values of all the phylogenetic distances between all organisms in a phylum. To analyze the relationship between scaling exponents and phylogenetic distances, we used the absolute values of the differences between the scaling exponents of the respective phyla as well as the exponent quotients.

## RESULTS AND DISCUSSION

### Scaling Shifts with Genome Size

Many scaling theories predict asymptotic behavior at the small and large end of a group, where certain properties go to zero or infinity. For example, growth rates and ribosome content all have asymptotes at either the small or large end of bacteria and/or eukaryotes (9, 10, 29). These physiological asymptotes appear in bacteria, microbial eukaryotes, mammals, plants, and insects (29) and can be seen as curvature away from the scaling relationships that capture the intermediate cell sizes. Thus, we should expect curvature in protein function scaling laws as most aspects of bacterial physiology are defined by scaling laws with asymptotes at the boundaries (9, 10, 14, 29). To search for this in a general manner, we performed a breakpoint analysis to see when there is support for a bilinear fit.

We discuss scaling between the COG categories, designating linear scaling or isometric growth for scaling exponents near one, sublinear or fractional dilution for exponents less than one, and superlinear or fractional enrichment for scaling exponents more than one. A high scaling exponent means more unique proteins are added and does not indicate the total number of proteins, for we do not address copy number in our study.

Plotting the number of COG category annotations against total protein annotations shows that, for the case of bacteria and archaea, the best fits for some categories are by more than one line (Figure 1 and Table 1). Such scaling shifts may be present in Eukaryotes but are not observed in our current work, likely due to the small number of genomes available.

**Figure 1:**
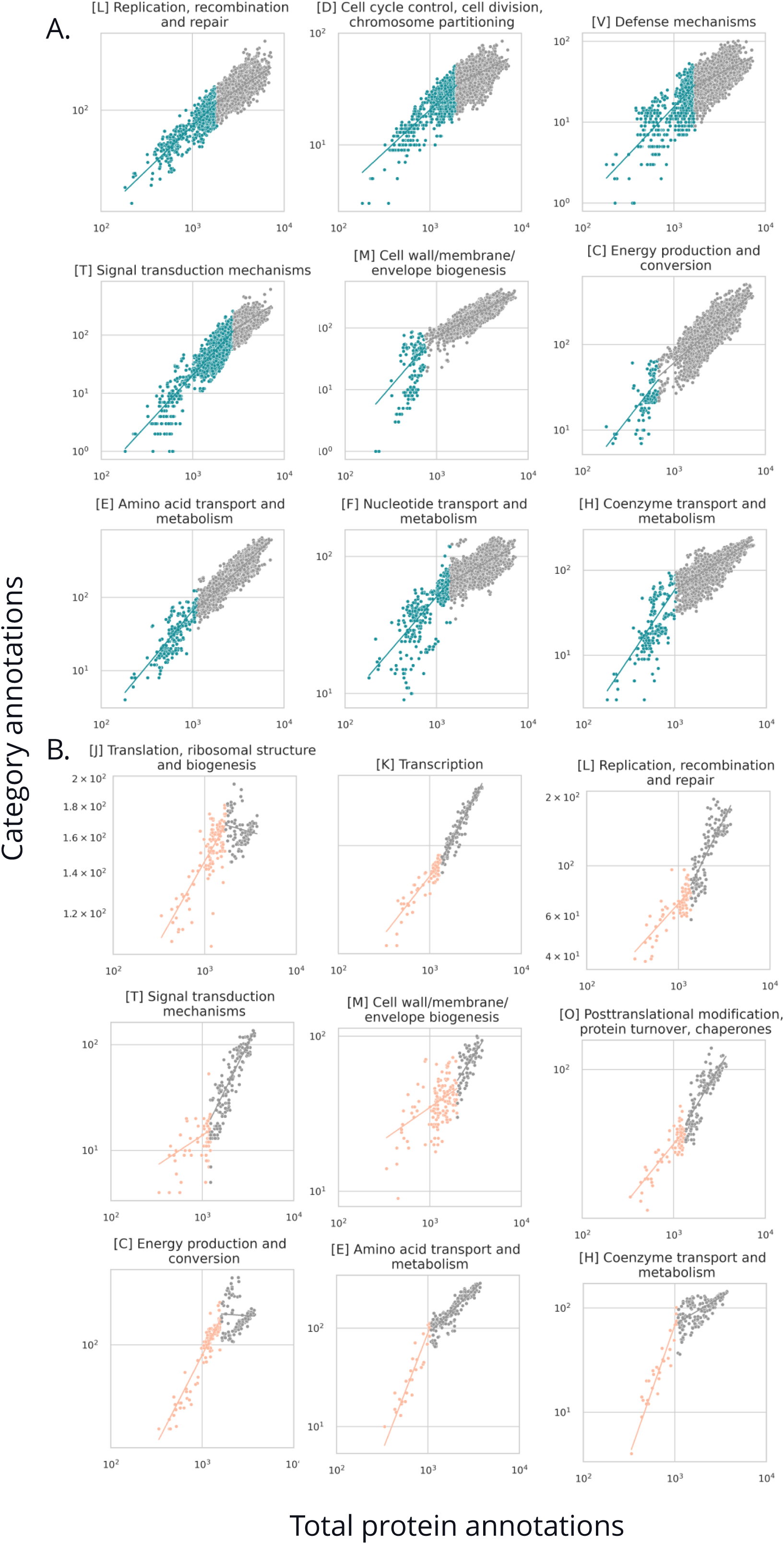
Breakpoint plots in A. Bacteria and B. Archaea. Axes are in log scale. The two different colors signify two slopes divided by a breakpoint.

**Table 1:**
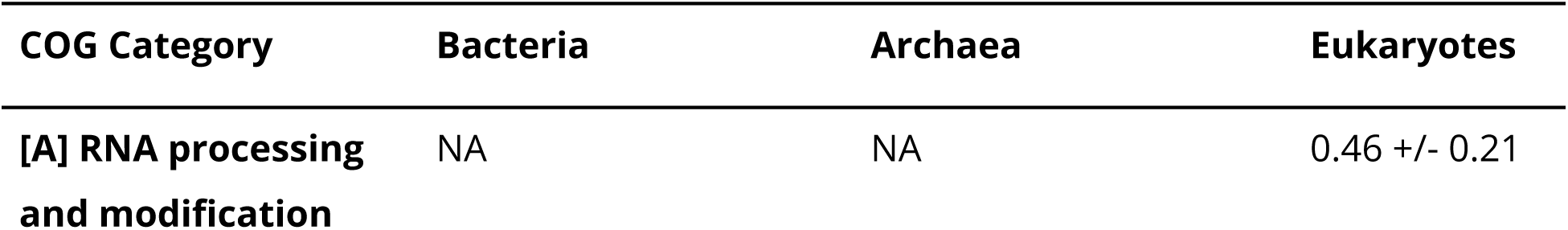
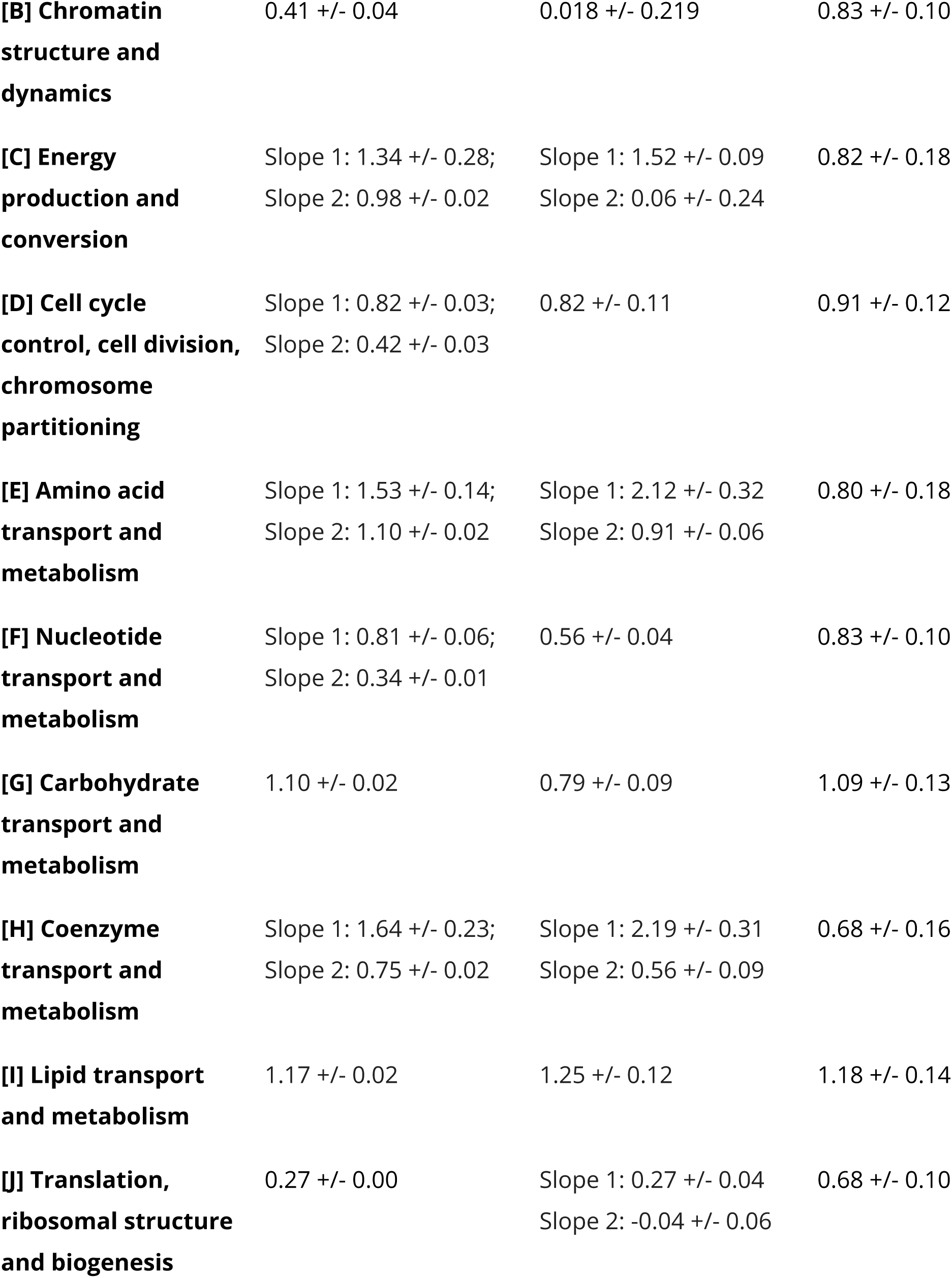
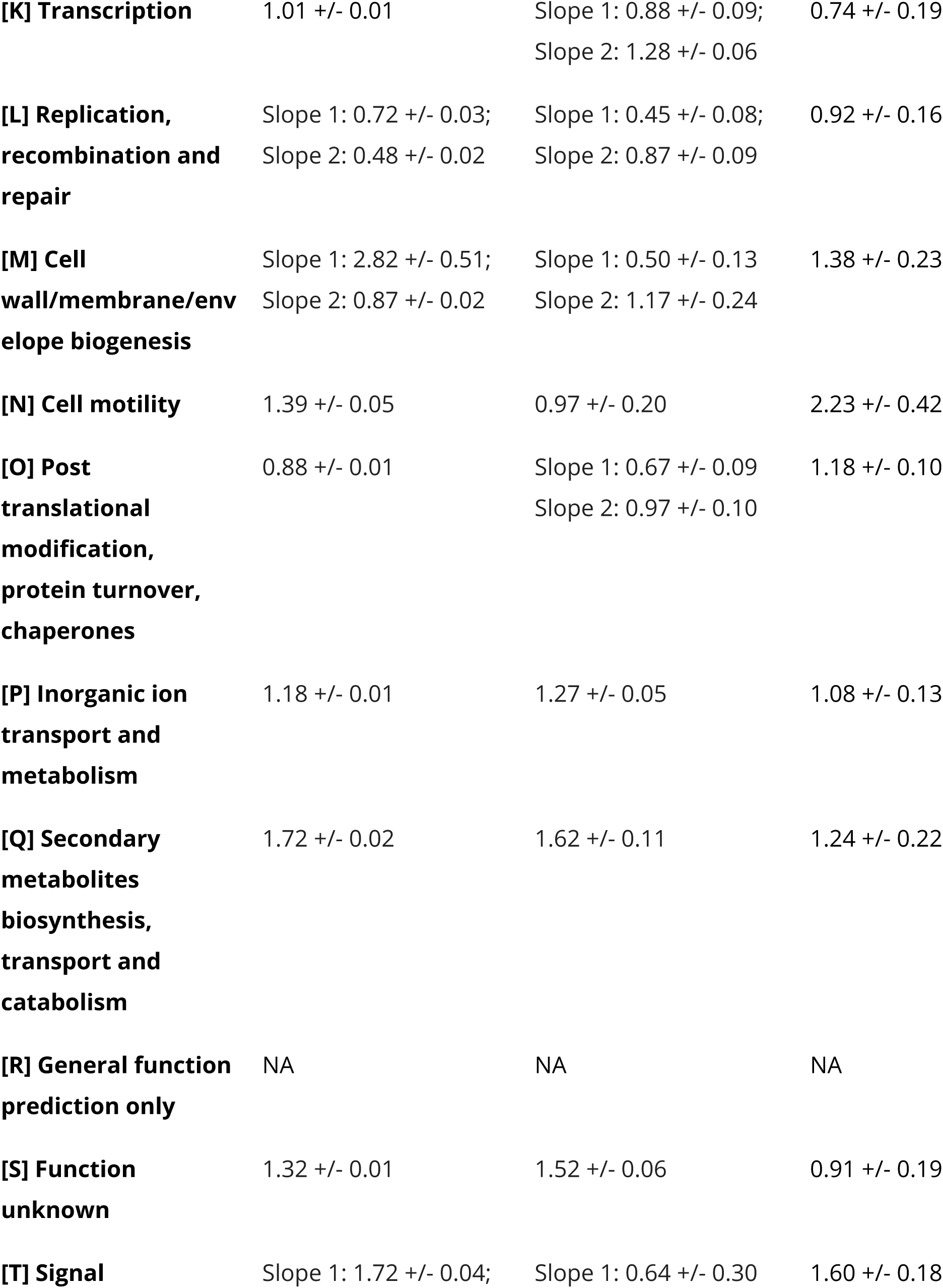
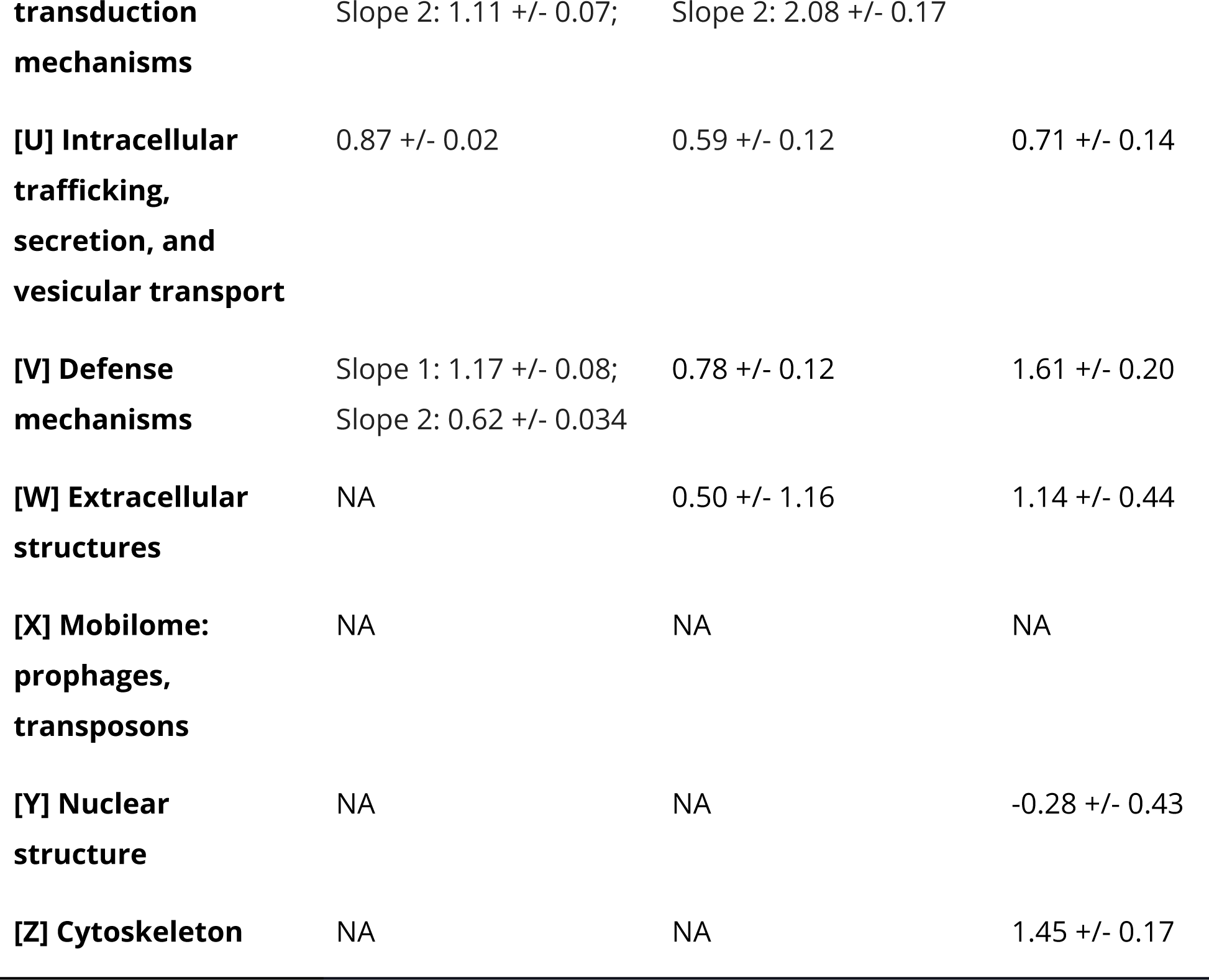
D**o**main **level scaling slopes with breakpoints.** Slopes 1 and 2 indicate categories with breakpoints. Supplementary Table 2 has Asgard archaea, DPANN and CPR separated from archaea and bacteria respectively.

Between archaea and bacteria, scaling shifts can be unique (Figure 1 and Table 1). The categories: [L] replication, [T] Signal transduction mechanism, [M] cell wall/membrane/envelope biogenesis, [C] Energy production and conversion, [E] Amino acid transport and metabolism, and [H] Coenzyme transport and metabolism are found in both bacteria and archaea, while [J] translation [K] transcription, and [O] are unique to the archaea. All scaling shifts in bacteria are all toward less positive exponents as the total number of annotations increases, however, in archaea, most shifts are to higher exponents except for [C], [E], and [H], which decrease as with bacteria. Categories [D] cell cycle control, [V] Defense mechanisms, and [F] nucleotide transport and metabolism are unique to displaying scaling shifts with bacteria.

### Domain Level of Functional Scaling

Next, we sought to gain an overview of functional scaling between the archaeal, bacterial, and eukaryotic domains as it occurs between different protein functional classes, looking at the broad groups of the clusters of orthologous groups.

#### Information Storage and Processing: COG categories A, B, J, K, and L

Ribosomes have been under strong selection pressures and are vital in differentiating the three domains of life. Interestingly, specific ribosomal proteins have been progressively accumulated and/or lost through evolution in the three domains (31). As has been reported by previous scaling studies (12, 13, 15, 25), we observed sublinear scaling for category **[J] Translation, ribosomal structure, and biogenesis** across all three domains. Archaea show an extreme reductive evolutionary scenario (Table 1), and archaea have been shown to preserve ribosomal proteins differently than bacteria and eukaryotes, losing up to ten ribosomal proteins in contrast to only four being lost in bacteria and eukaryotes through evolution (31). Eukaryotes scale with a mildly sublinear scaling exponent of 0.68 +/- 0.10. The normalization constants for this category are exceptionally high for all the domains, which indicates large quantities of proteins in this category from the onset of genome evolution as we can see it and explains ribosomal proteins involved in several extra-ribosomal functions, mainly in transcription and regulation (Supplementary Table 7 and 8) (32). Since in extant life, ribosomes are almost 2/3rd rRNA, and only 1/3rd proteins, a deeper understanding of rRNA evolution would help draw answers to the reductive co-evolution of ribosomal proteins (33).

Since many signal transduction pathways are heavily dependent on transcription factors, it is essential to note that the COG system of annotation separates transcription factors and proteins involved in signal transduction. While some proteins have both **[K] Transcription** and **[T] Signal transduction mechanism** annotations, in our study, we have separated these annotations and counted them individually for an organism (for example, a protein annotated as KT is counted once as K and once as T). Similar to the observations using COG annotations by Cordero et al. (23), we did not observe quadratic scaling for the category **[K] Transcription,** which has been previously reported by studies that used the GO-PFAM annotation (13, 34). We observed near-linear scaling in archaea, and bacteria and lenient sublinear scaling in eukaryotes. Complex transcription regulation mechanisms might explain the sublinear scaling in eukaryotes, as complexity often gives rise to more integrity in connective regulation, where proteins participate in more functions (34). A more detailed analysis of the role of non-coding DNA and chromatin in transcription regulation and a more comprehensive sample size of eukaryotic species would help us better understand scaling trends of transcription regulation in eukaryotes. We observe superlinear scaling in signal transduction, as discussed ahead.

DNA replication allows organisms to pass on information, mutate, evolve, and adapt through evolution. Proteins in this category are required by the cell at specific time points in the lifecycle of a cell. They are well known to be similar between eukaryotes and archaea but different in bacteria (35). We highlight similarities and differences in their scaling patterns that have not previously been observed (36, 37). Eukaryotes show almost linear scaling (0.92 +/- 0.16) with genes involved in **Replication, recombination, and repair [L],** while archaea show sublinear scaling, with the dilution being greater in smaller organisms (Table 1 and Figure 2). Eukaryotes are believed to have not added unique subunits to the replisome machinery, and the subunits remain the same in the modern replisome machinery as the ancestral machinery (36). While we agree on eukaryotes, we differ from (36) in reporting that archaea scale sublinearly and do not show a continually increasing trend of unique subunits. Instead, the best fit is achieved by two lines of slope less than one, with smaller genomes scaling at 0.45 +/- 0.08 and larger genomes being fit with an exponent of 0.87 +/- 0.08. Additionally, we observed that bacteria support a breakpoint at about 1/4th of the total protein annotations. The initial 25% of the data points show a sublinear scaling exponent of 0.72 +/- 0.03, while the later data points scale even slower with an exponent of 0.48 +/- 0.02. (Table 1), opposite the trend in archaea where the exponent increased.

**Figure 2:**
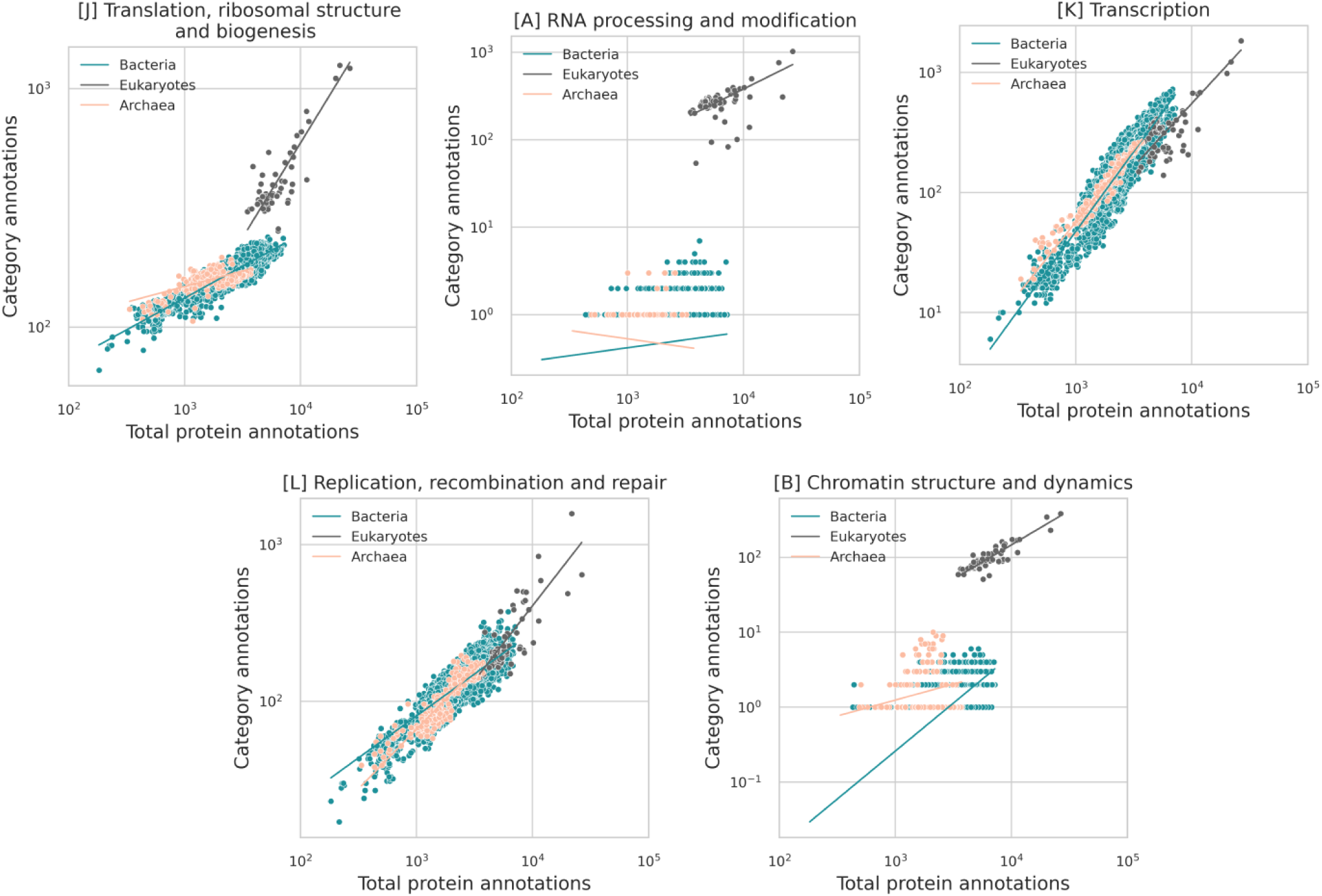
Domain level scaling in protein classes associated with information storage and processing. Axes are in log scale.

**Figure 3:**
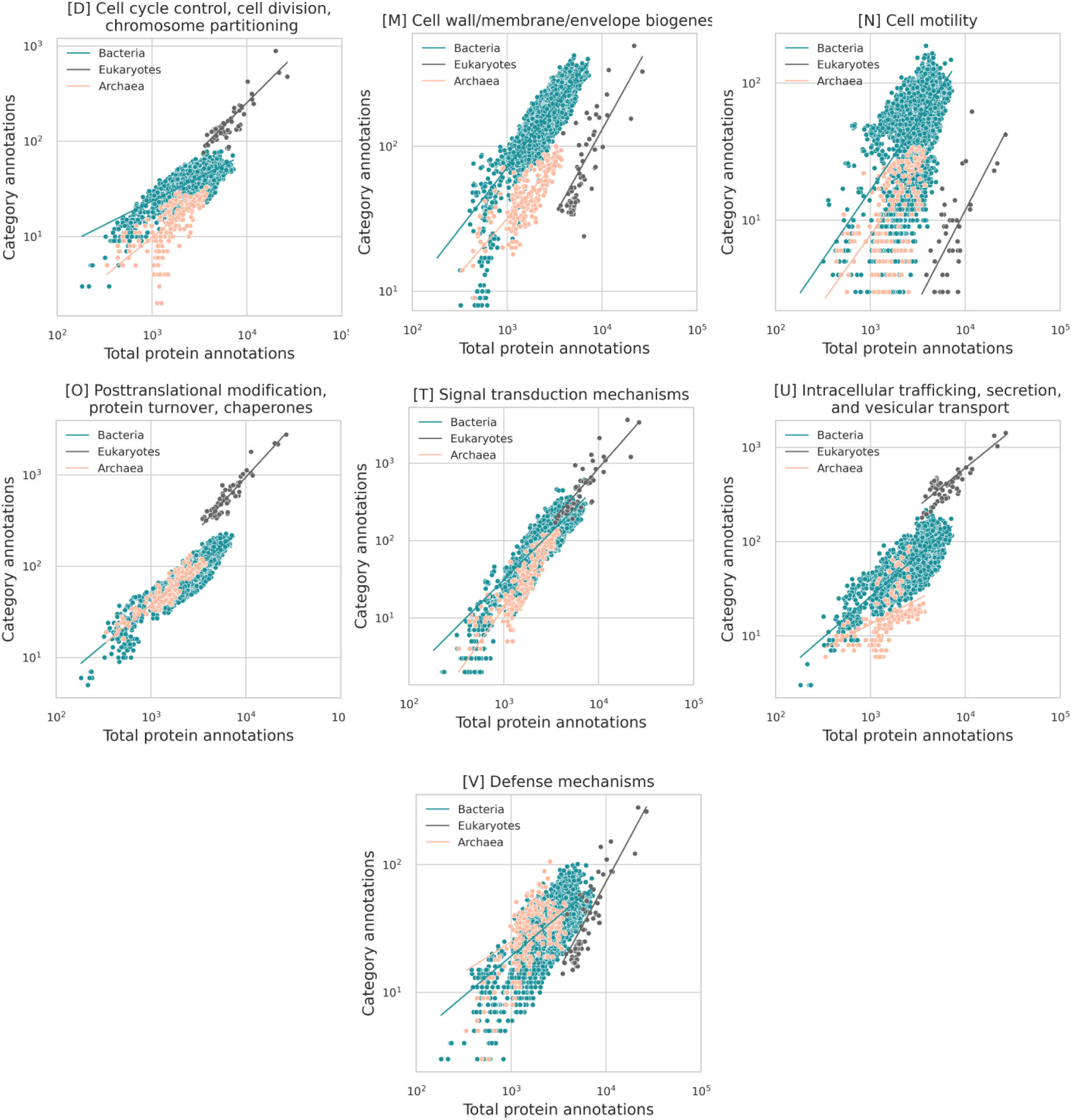
Domain level scaling in protein classes associated with cellular processing and signaling. Axes are in log scale.

Although archaea do not show a trend in **[B] Chromatin structure and dynamics**, Asgard archaea, when looked at alone, do (1.24 +/- 0.70), similar to what is seen in eukaryotes (0.83 +/- 0.10), while bacteria show sublinear scaling with an exponent of 0.41 +/- 0.04 (Table 1 and Supplementary Table 1). eggNOG includes annotations of proteins that carry out histone acetylation, acetoin utilization, certain chromatin remodeling proteins, and other kinase activity proteins in bacteria. Although other nucleoid-associated proteins are not explicitly annotated (38), a more detailed analysis of bacterial chromatin is needed.

The spliceosome, considered among the most complex modern cell machineries, is believed to have evolved its complexity through neutral evolution (39, 40). Despite this, we observe a sublinear scaling trend for **[A] RNA processing and modification** in eukaryotes, indicating that the spliceosome machinery complexity does not become enriched in the 79 unicellular eukaryotes we considered (Table 1). The much discussed constructive neutral evolution in RNA processing enzymes (41–43) would be a future target where genomes would be sampled from LECA across more complex eukaryotes to identify where scaling increased.

#### Cellular Processes and Signaling: COG categories D, M, N, O, T, U, V, W, Y, and Z

Previously, van Nimwegen observed sublinear scaling for cell cycle proteins (12). In bacteria, **category [D] “Cell cycle control, cell division, chromosome partitioning”** data supports a breakpoint from about near linear scaling (0.82 +/- 0.03) below 30% of the bacterial genome sizes and is sublinear (0.42 +/- 0.03) after that (Table 1, Supplementary Table 3). We observe near-linear scaling for archaea and eukaryotes (Table 1).

Membrane proteins in prokaryotes encompass about 20-30% of the proteins expressed, and an evolutionary leap from simple to complex cells would have meant increasing the number of membrane proteins owing to the increase in special membrane features and newer compartments in cells (44). Proteins in category **[M] “Cell wall/membrane/envelope biogenesis”** scale superlinearly for small bacterial genomes (2.82 +/- 0.51) while showing a mild sublinear scaling after about only 10% of total protein annotations on the X-axis (0.87 +/- 0.02) (Supplementary Table 3). Apparently, as genome size increases from the smallest end, more unique proteins are required, whereas less diversity is required towards the larger scale. The breakpoint analysis in archaea reveals the opposite trend, with the exponent increasing with the total number of annotations from 0.05 +/- 0.13 with a high scatter to 1.17 +/- 0.24. In eukaryotes, the scaling is superlinear (1.38 +/- 0.23), and, interestingly, scaling after the breakpoint in archaea is similar to this value as endosymbiotic theories support the idea that Asgard archaea engulfed bacteria to form the eukaryotes we know today (44).

In relation to a transition from Asgard to Eukarya, we observed a nearly linear scaling in **[U] Intracellular trafficking, secretion, and vesicular transport proteins** for Asgard archaea (1.17 +/- 0.53) (Supplementary Table 1). The endomembrane system that consists of the Golgi apparatus, endoplasmic reticulum, vacuoles, and vesicle traffic is conserved in all sizes of eukaryotes, indicating their conserved functioning from the Last Eukaryotic Common ancestor (LECA) (44), agreeing with our sublinear scaling exponents for category U in eukaryotes (0.71 +/- 0.14).

It is also interesting to observe that while category [M] supports a breakpoint for bacteria, it does not for category [U], hinting towards the universal scaling pattern for intracellular trafficking, secretion, and vesicular transport proteins across a range of bacterial cell sizes (Supplementary Table 3). More exploration into specific scaling patterns of these two categories could unfold the mechanisms behind the evolutionary scaling of proteins involved in building the cell wall membrane/ envelope biogenesis systems and the endosymbiotic theory for the origin of eukaryotes from mitochondria.

**[T] Signal transduction mechanisms** scale superlinearly in archaea after a breakpoint, changing from 0.64 +/- 0.30 to 2.08+/- 0.17. In eukaryotes, the scaling is also superlinear, whereas in bacteria, the slope is 1.72 +/- 0.04 for bacteria below 40% of the total annotations on the X-axis, and the trend takes a linear shift (1.11 +/- 0.07) for the larger organisms in bacteria (25) (Supplementary Table 3). These observations differ from previous studies, which show a continuous trend in the scaling of signal transduction mechanisms and for bacteria hint at increased efficiency through evolution, with cells using the same unique subunits to create more complicated machineries for more complex organisms as evinced by the near linear exponent after the breakpoint. For archaea, however, the case is very different, and category [T] is enriched during genome expansion.

We observed superlinear scaling in eukaryotes for proteins in category **[V] Defense mechanisms** (1.61 +/- 0.20) (Table 1). For bacteria, we observe an exponent of 1.17 +/- 0.08 for organisms up to about 23% of the total protein annotations (Supplementary Table 3). The scaling slows down to 0.616 +/- 0.034 after that, and in archaea, the scaling is sublinear (0.78 +/- 0.12). Defense genes in prokaryotes tend to cluster into “defense islands,” which are highly susceptible to gene shuffling and rapid evolution, and many of these islands’ genes are uncharacterized, hinting that a greater inspection of these clusters would give us a better overview of the evolution of prokaryotic defense strategies (45).

In Eukaryotes, the splicing machinery and defense mechanisms are often related (46, 47). Tracing this back to LECA, it is believed that the group II retroelements from the proteobacterial cell infected the host archaeal genome (46, 47). The host developed defense mechanisms against these introns; some were retained, some were lost, and some were newly developed (47). What Koonin describes as an “intron catastrophe” could have led to an increase in processes like ubiquitin signaling acting as defense mechanisms: previously offering different functions, these mechanisms were already present in early cells which explored these mechanisms temporarily to defend against external invading nucleic acids (40). Constructive neutral evolution events might also answer why random events like ubiquitination become fixed over evolution as one of the mechanisms under the big umbrella of defense mechanisms (40).

Environmental surfaces would have influenced the evolution of bacterial motility, and in contrast, cell enlargement must have been an influencing factor for eukaryotes (48). We observe steep superlinear scaling in **[N] Cell motility** for eukaryotes and Asgard archaea with scaling exponents above two, while bacteria and archaea show scaling exponents close to one (Table 1 and Supplementary Table 1).

Through evolution, organisms must have increased their protein turnover rate to compensate for mistranslation and maintain proteome homeostasis (49). We report an almost linear scaling for proteins involved in **[O] Post-translational modification, protein turnover, and chaperones** for all domains. Why we observe linear and superlinear scaling in this category could be due to the same process happening faster without the need to recruit new proteins.

#### Metabolism: COG Categories C, E, F, G, H, I, P, and Q

The relationship between metabolism and cell size has been extensively discussed in several works (9, 10, 14, 17, 50). It has previously been observed that categories corresponding to metabolism scale almost isometrically with genome size, with metabolic networks tending to expand proportionally with genome size (25). We highlight the different scaling patterns observed here.

**[C] Energy production and conversion** supports a breakpoint for bacteria very early (before 10% of the total protein annotations) with a similar trend as observed with all the breakpoints in bacteria in our analysis (Supplementary Table 3). The scaling is superlinear (1.34 +/- 0.28), then slows to near-linear (0.98 +/- 0.02). Archaea scale superlinearly until 42% of total protein annotations (1.52 +/- 0.09). (Supplementary Table 4). In archaea, category [C] is notable since the data was split into two lines as the number of annotations increases (this is not captured in the slopes given in table 1 due to the way that the regression was done). Eukaryotes also show near-linear scaling in [C] (0.82 +/- 0.18).

**[E] Amino acid transport and metabolism** show superlinear scaling for smaller bacterial genomes (1.53 +/- 0.14) and near linear scaling for the larger ones (1.10 +/- 0.02). Archaea show superlinear scaling for the first 28% of the total protein annotations, but the scaling falls to linear after that (Supplementary Table 4), and for Eukaryotes, the scaling is just below linear. **[F] Nucleotide transport and metabolism** is sublinear across the domains, with a breakpoint detectable in bacteria. Perhaps this progressive delusion is related to very early nucleotide metabolism in the LUCA. **[H] Coenzyme transport and metabolism** shows superlinear scaling for both smaller bacteria and archaea genomes and sublinear scaling for bacteria and archaea with larger genomes, whereas eukaryotes are sublinear. Category **[I] Lipid transport and metabolism** is nearly linear for all three domains (Supplementary Table 3).

**[P] Inorganic ion transport and metabolism** shows similar scaling trends in both the domains, and **[Q] Secondary metabolites biosynthesis, transport, and catabolism** is superlinear for all domains but less steep in eukaryotes than bacteria and archaea.

### Variability between microbial phyla

The observation that scaling relationships were best fit in many cases in bacteria by multiple lines led us to hypothesize that phyla specific scaling relationships exist within the

domains. Figure 5 shows the spread of scaling between microbial phyla for a number of COG categories (all of the COG categories are shown in Supplementary figure 4). The figure also includes microbial Eukarya, which were not taxonomically separated due to fewer genomes.

Across the three domains, the ribosome is enriched in subunits from bacteria to archaea to eukaryotes which have the highest number of subunits (31). **Category [J] Translation, ribosomal structure, and biogenesis** contains these proteins and others and shows trends of phyla-specific reductive evolution and some expansion. Eukaryotes show the closest exponent to one, and Asgard archaea fractionally dilute less compared to other archaea (0.36 +/- 0.15) as compared to other archaea (0.14 +/- 0.02) (Supplementary Table 1). In microbes, ribosome usage increases with genome size due to proportionality between size and growth rate (9, 10), but the functional scaling observed is primarily sublinear, indicating that the usage is derived by using more of the same thing and not an elaboration of more types.

The scaling of regulation has received significant attention in past studies (13, 16, 25, 51). We observe phyla-specific scaling in categories [K] and [T]. Alpha, Beta, Gamma, Delta, and Epsilonproteobacteria show scaling with an exponent of around 1.5 (Supplementary Table 2). Additionally, Planctomycetes, Spirochaetes, and Verrucomicrobia show high scaling exponents in addition to the members of Proteobacteria. All other phyla show exponents below 1.4 (Supplementary Table 2 and Supplementary Figure 4).

Deviating from the nearly universal trend of superlinear scaling in **[T] Signal transduction mechanisms** are Chlamydiae and Spirochaetes, two groups notoriously known to be pathogenic, in addition to Acidobacteria (Supplementary Table 2 and Supplementary Figure 4). These three groups show linear scaling. Crenarchaeota in archaea show sublinear scaling (Supplementary Table 2 and Supplementary Figure 4). This is particularly interesting to note because Crenarchaeota use only one type of phosphorylation (Hanks type phosphorylation) and not two-component signal transduction like Euryarhcaeaota (52).

DPANN and CPR show superlinear scaling for **[F] nucleotide and [E] amino acid transport and metabolism** (Figure 5, Supplementary Table 2, and Supplementary Figure 4). Figure 5 for [F] is interesting to observe. As CPR and DPANN increase in unique protein annotations per genome, this functional category might be among the first additions they scale rather rapidly compared to the scaling of other proteins. DPANN shows an exceptionally high exponent for **[H] Coenzyme transport and metabolism,** while CPR has an exceptionally high exponent in **[I] Lipid transport and metabolism** (Supplementary Table 2). No known lipid biosynthesis pathways are known for CPR (53). Moreover, due to their exceptionally small sizes, CPR membranes could experience stress due to the high curvature to area ratio, explaining the abundant presence of lysolipids (54). CPR might also receive or forage certain lipids from surrounding bacteria in communities (54). Although the reason behind high scaling exponent, which would mean that as the size of bacteria in this group increases, more lysolipids (which contradicts the high curvature to area ratio point) or perhaps scavenged lipids increase, remains to be understood. These data make it intriguing to consider that there are multiple paths into, or out of, a reduced genome size state.

Tenericutes that lack a peptidoglycan cell wall, and Chloroflexi, which have a highly folded membrane but are somewhat enigmatic in terms of their membrane composition (55, 56), showed quadratic scaling of proteins involved in **[M] cell wall, membrane, and envelope biogenesis** (57, 58). Tenericutes also drive the drastic superlinear scaling in the initial part of the bacterial domain scaling plot (Figure 8). DPANN are superlinear in [M], again contrasting to CPR, which are sublinear.

**[Q] Secondary metabolites biosynthesis, transport, and catabolism** show universal superlinear scaling trends except for three groups which also have a wide confidence interval (Supplementary Table 2). **[N] Cell motility** is highly variable across microbial phyla. Although Eukaryotes and Asgard are still amongst the highest scaling exponents, there are other groups that stand out from their cumulative domain level scaling exponents, like Cyanobacteria, Verrucomicrobia, Synergistetes, and Firmicutes (Supplementary Figure 4 and Table 2).

Planctomycetes show some of the slowest scaling exponents in **[D] Cell cycle control, cell division, chromosome partitioning** (Supplementary Table 2 and Supplementary Figure 4). Planctomycetes lack the multi-complex divisome most bacteria use for binary fission (59). While the budding mechanism in bacteria of this phyla is not well understood, they use an extremely conserved component of the divisome (59).

Considering the expanded prokaryotic taxonomy considered in this study, the prominently varying scaling laws may highlight the differences in the physiology and architecture of prokaryotes. To quantify the deviation of the individual phyla exponents from the scaling exponents observed at the domain level, we calculated Z-statistic scores as before (13). Figure 6 arranges the COG categories by the range of the z-statistics scores in each category which were calculated for bacteria and archaea separately. These summarise how individual phyla/clade exponents differ from their respective domains (archaea and bacteria).

First, we can see that archaea and bacteria are unique in how variation is distributed across functional categories (Figure 6 and Supplementary Table 6). For bacteria, **[H] Coenzyme transport and metabolism** showed the most variation (Figure 6 and Supplementary Table 6). The smallest variation was observed in **[B] Chromatin structure, [Q] Secondary metabolites biosynthesis, transport and catabolism** and dynamics, and **[V] Defense mechanisms**, indicating that the bacterial domain exponents are a good representative of the microbial diversity for these categories. Furthermore, Actinobacteria are outliers. Cyanobacteria show extreme trends too. Gemmatimonadetes primarily lie near zero, indicating exponent values similar to the whole domain. Although these scores allow us to summarize the variability or similarity in exponents, it is important to note that the Z-score does not include the sample size of the groups, and there is no means of identifying the confidence intervals.

Amongst archaeal phyla, Euryarchaeota are significant outliers (Supplementary Table 6). We hypothesize that this could be explained by “slow scaling hypothesis,” which considers that as the genome size increases, there is no need for multiple copies of a protein or metabolic diversity. Thaumarchaeota lies mainly near zero (Supplementary Table 6). As with bacteria, **[H] Coenzyme transport and metabolism** shows the most significant variation indicating the different physiological and functional organization of this category across phyla. Lineages differ with respect to their functional repertoires (60), and we have demonstrated in this section how variable scaling is between phyla. These observations follow van Nimwegen et al. in showing that functional scaling is also unique to particular groups.

Interestingly, when looking at Figures 4 and 6, we observe that metabolism is the most similar between domains (data clouds overlapping on top of each other) but some of the most variable categories between phyla (from Z-scores).

**Figure 4:**
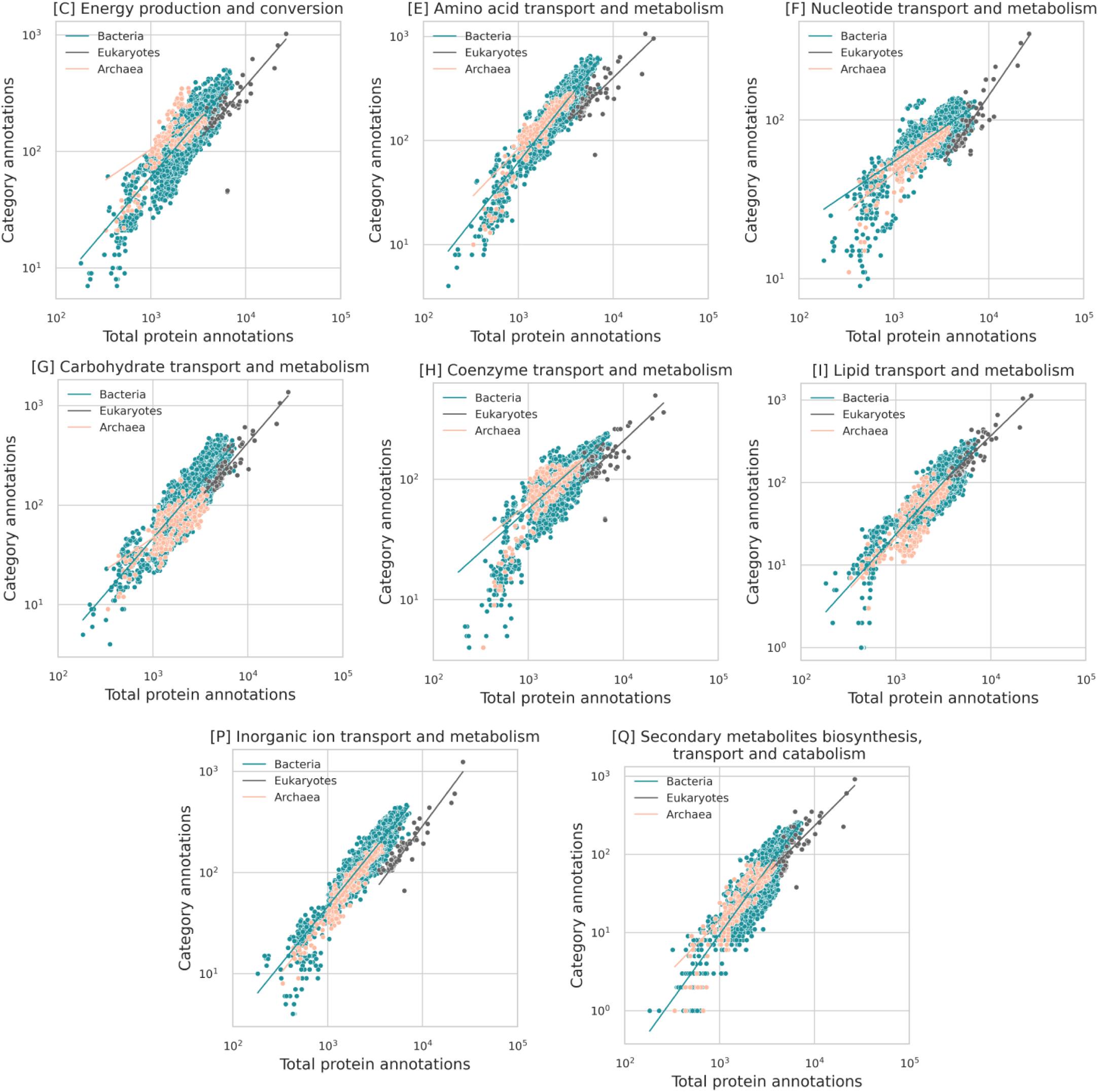
Domain level scaling in protein classes associated with metabolism. Axes are in log scale.

**Figure 5:**
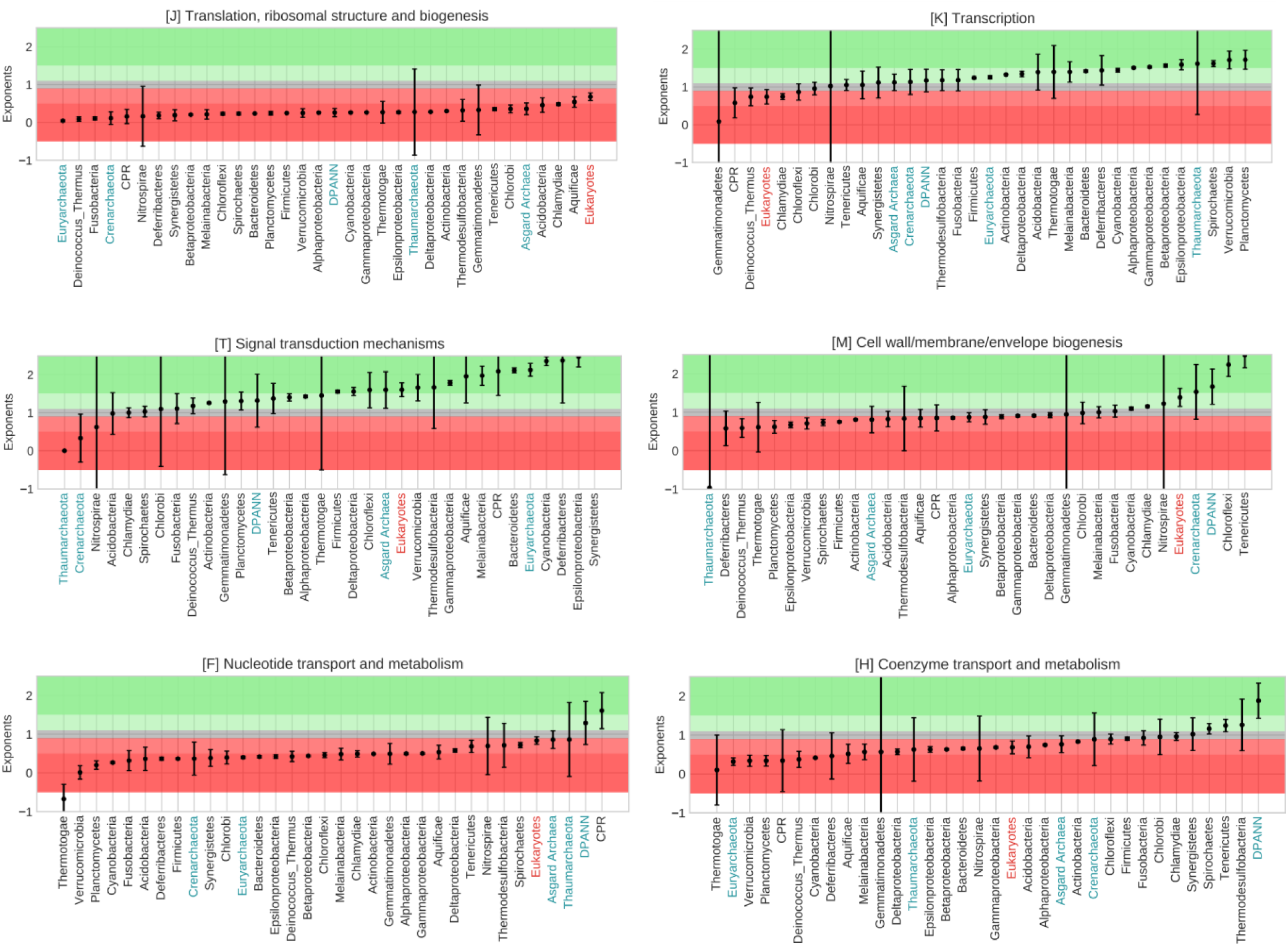
Phyla arranged in the increasing order of their exponents with 95% confidence intervals for selected COG categories. Background horizontal span colors signify scaling. From dark green to dark red, meaning superlinear scaling to sublinear scaling. The grey span in between signifies linear scaling. A full figure with all COG categories is available in the supplementary material.

**Figure 6:**
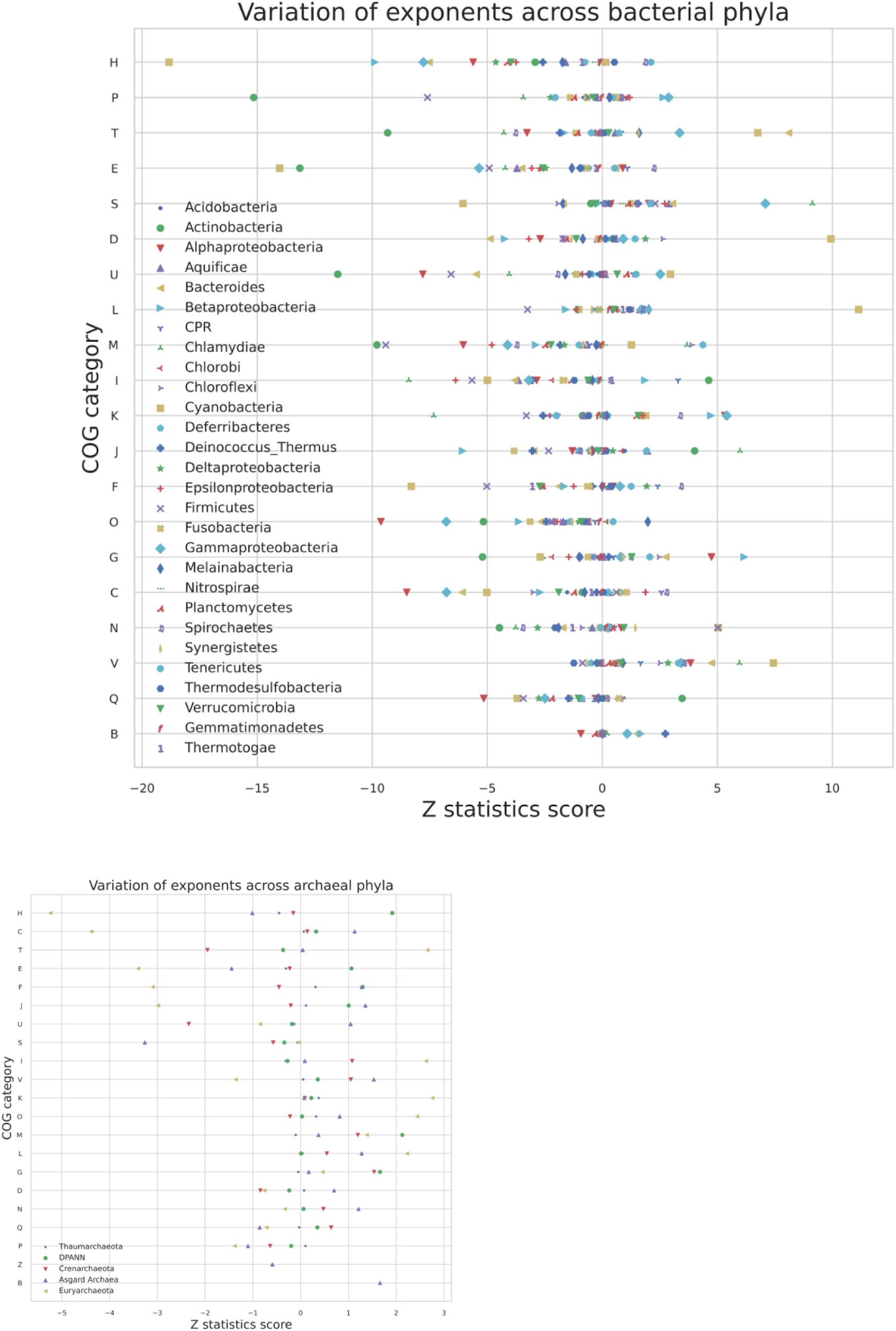
Z-statistics plots showing the difference between scaling exponents of individual phyla from their respective domains.

**Figure 7:**
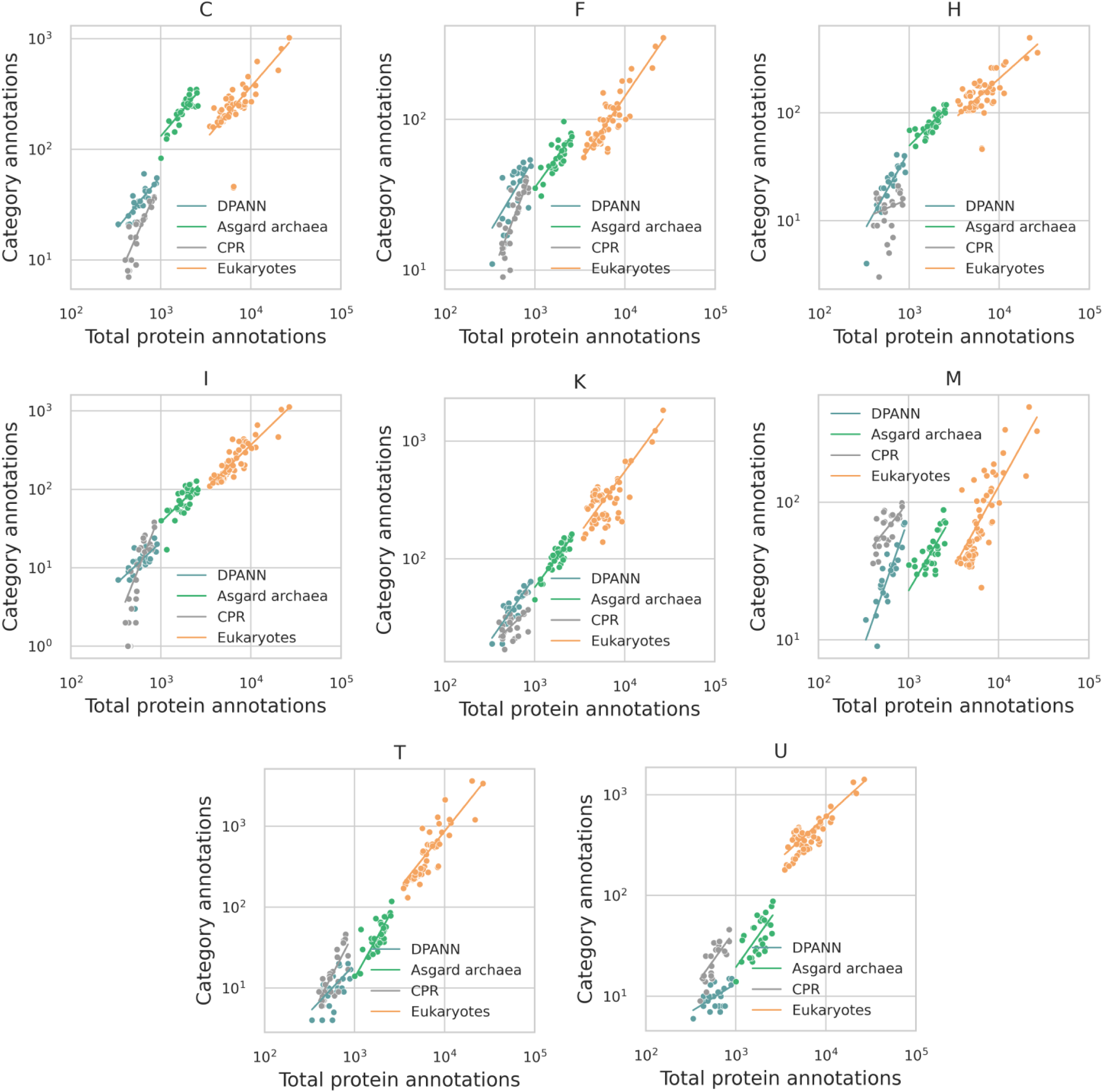
Scaling comparison between CPR, DPANN, Asgard archaea, and Eukaryotes

### Evolutionary Edges: Asgard Archaea, DPANN, CPR

Organisms with small genomes like DPANN and CPR that lack major metabolic pathways show markedly high scaling exponents for proteins belonging to nucleotide and amino acid transport and metabolism (53) (Figure 5, Supplementary Figure 4, Supplementary Table 2). This could be explained by the vast variety of cellular activity that the proteins in these categories carry out in contrast to categories of the central dogma of biology with rather specific roles. It is interesting to note how CPR shows sublinear scaling in category [H] Coenzyme transport and metabolism, while DPANN shows almost quadratic scaling in this category (Figure 5 and Supplementary Table 2), suggesting some possible major differences between functional repertoire adjustment during genome expansion or shrinkage in these two very different phylogenetic groups. In addition to [H], CPR, and DPANN show different scaling patterns in [C] Energy production and conversion, [I] Lipid transport and metabolism, [K] Transcription, [M] Cell wall/membrane/envelope biogenesis, [T] Signal transduction mechanisms, and [U] Intracellular trafficking, secretion, and vesicular transport while being similar in the rest of the categories.

These patterns are shown for a subset of COG categories in Figure 7, and for all the COG categories in Supplementary Figure 5. Asgard archaea and eukaryotes have similar scaling exponents for all categories except [J] and [K] (Table 2, Supplementary Table 2, and Supplementary Figure 5).

**Table 2:**
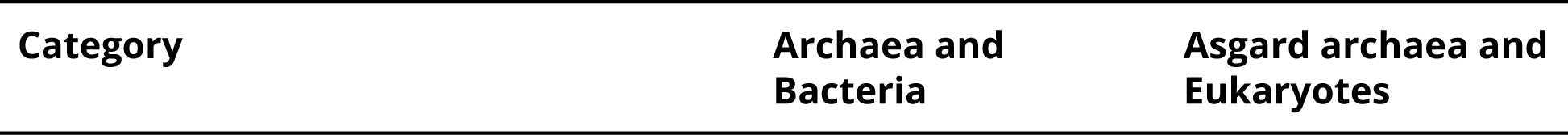
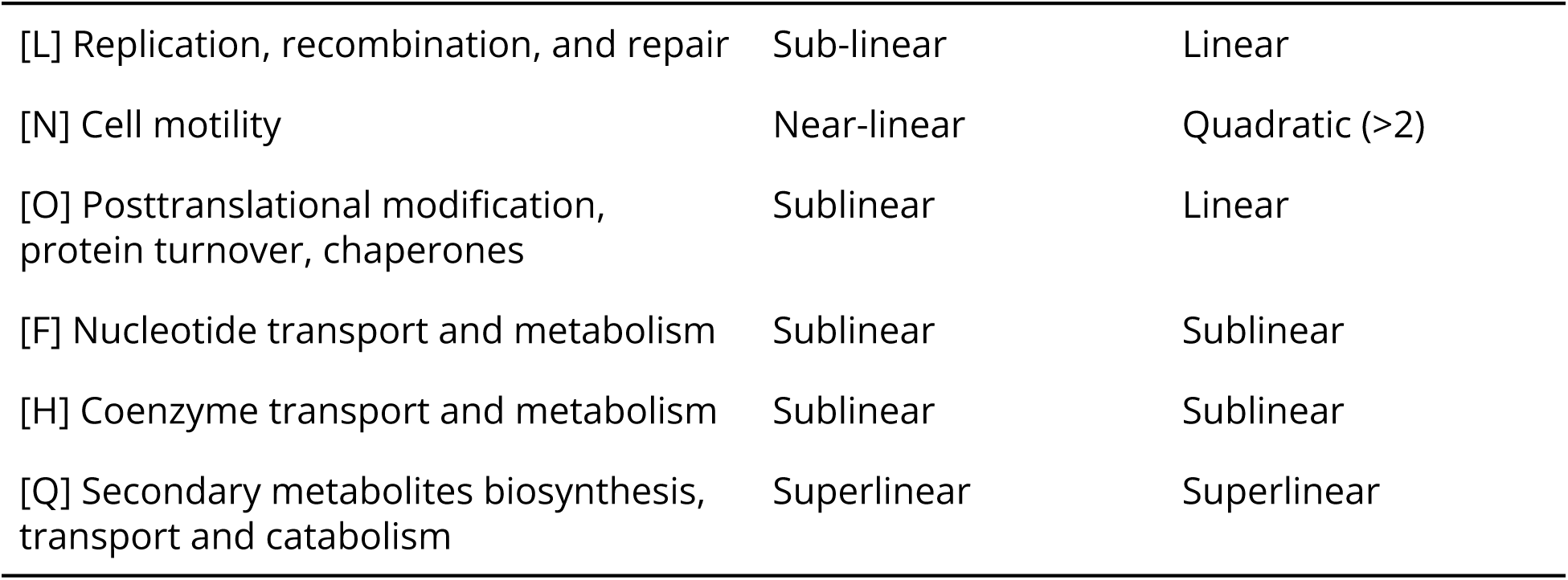
Similarities and differences between functional scaling within archaea and bacteria in comparison to Asgard archaea and Eukaryotes.

| Category | Archaea and Bacteria | Asgard archaea and Eukaryotes |
| --- | --- | --- |
| [L] Replication, recombination, and repair | Sub-linear | Linear |
| [N] Cell motility | Near-linear | Quadratic ($>2$) |
| [O] Posttranslational modification, protein turnover, chaperones | Sublinear | Linear |
| [F] Nucleotide transport and metabolism | Sublinear | Sublinear |
| [H] Coenzyme transport and metabolism | Sublinear | Sublinear |
| [Q] Secondary metabolites biosynthesis, transport and catabolism | Superlinear | Superlinear |

### Variability of Scaling Within Microbial Phyla

We hypothesized that phyla-specific scaling might explain the observation of breakpoints analysis (Figure 1). Based on the median protein annotations, we might expect some COG categories to be crossed over more than others. However, when we overlaid the total protein annotations in each phylum with the breakpoint locations, we found that breakpoints are spanned by phyla and do not appear to be an evolutionary barrier to taxonomy (Figure 8).

**Figure 8:**
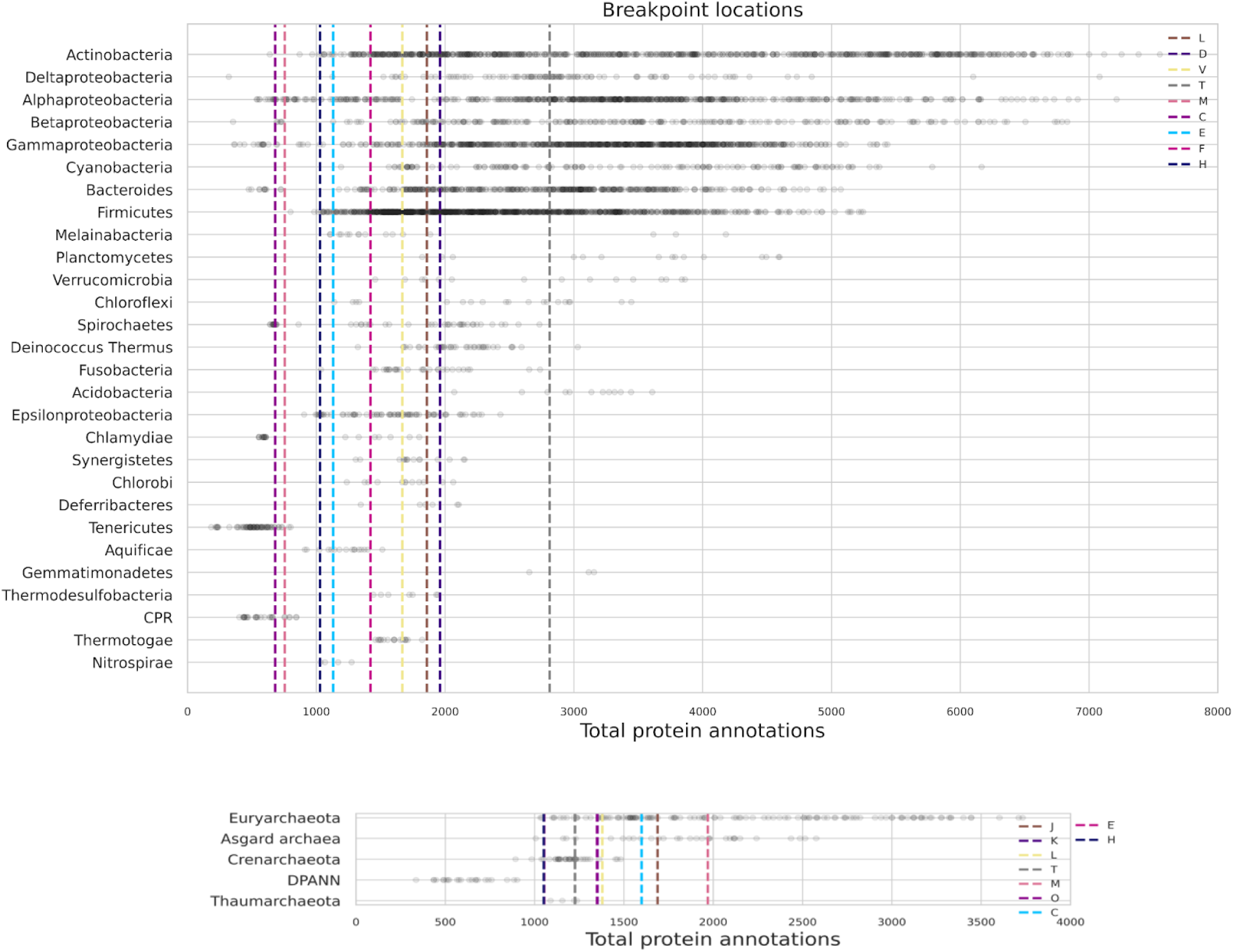
Breakpoint locations across bacterial and archaeal genomes: Individual data points are total protein annotations of organisms. Phyla ordered from the most spanning to the least spanning on the X-axis from top to bottom.

Both archaeal and bacterial annotations show a breakpoint in categories [C] Energy production and conversion, [E] Amino acid transport and metabolism, [H] Coenzyme transport and metabolism, [M] Cell wall/membrane/envelope biogenesis, [T] Signal transduction mechanisms, and [L] Replication, recombination and repair.

For the metabolism category [C], while bacteria show a breakpoint towards the extreme low end, archaea have this breakpoint positioned near the center. For [E] and [H], archaea and bacteria both have the breakpoints positioned around the same number of annotations. The trends in the scaling before and after the breakpoints, as discussed in detail in the *Domain level of functional scaling* section above, is similar in both bacteria and archaea, with the scaling slowing down from superlinear scaling after the breakpoint and the slowing down being more prominent in [C] and [H] for both domains.

For cellular processing and signaling categories [M] and [T], the breakpoint locations are different between archaea and bacteria with the breakpoint for [M] lying near the lower end for bacteria and the higher end for archaea. For [T], the breakpoint falls near the higher end for bacteria, while it is centered for archaea, crossing all phyla except for DPANN. The patterns in the scaling are very different too. While [M] and [T] both scale superlinearly before the breakpoint for bacteria, they scale sublinearly for archaea. After the breakpoint, these categories scale sublinearly for bacteria but superlinearly for [T] and near linearly for [M] archaea.

Most of the breakpoints occur near the smallest genome sizes. This makes sense since a variety of dramatic physiological and compositional shifts are known to occur for the smallest cells (9, 10, 14). For example, growth rates, ribosomal abundances, and the space available for macromolecules all have asymptotic behavior for the smallest cells (10). The smallest cells are running out of internal space for macromolecules and metabolism is increasingly being dedicated to maintenance purposes leaving little for growth. These dramatic shifts impose serious constraints that are likely to be accompanied by corresponding shifts in the genome scaling as genomes are reduced.

Breakpoints for categories [D] Cell cycle control, cell division, chromosome partitioning, [V] Defense mechanisms, [F] Nucleotide transport and metabolism in bacteria and [J] Translation, ribosomal structure and biogenesis, [K] Transcription, [O] Posttranslational modification, protein turnover, chaperones in archaea are unique for their domains. Although we consider statistically significant differences, we must interpret cautiously and propose future work where breakpoint location would be associated with statistical support. Understanding the evolutionary significance of breakpoints would need extensive analysis of specific phyla. However, these idiosyncratic patterns in the positioning of breakpoints from a domain point of view hint at some evolutionary peculiar events.

### Genome scaling is not correlated with the phylogenetic distance between phyla

Within a species-level group of bacteria, the phylogenetic distance was found to be related to genome composition (61); we hypothesized that variability in scaling might be related to phylogenetic distance between members across the tree of life, and to investigate this we used a tree of life constructed from ribosomal proteins (30). We calculated the patristic distance between clades to determine if scaling relationships among phyla are related to taxonomy evolution. No clear evolutionary relationship between phyla and groups was observed (Supplementary Figure 3), suggesting that variability in scaling is not related to the evolutionary relationships between taxonomic clades, but instead may be related to environmental, physiological, or other variables.

### What do our results tell us about the earliest genomes?

Understanding universality or variability in scaling patterns across diverse species would help us work our way backward to understand what to expect in early evolutionary periods, where cells similar to those today may have emerged from smaller genomes. Due to an inability to copy large amounts of information with high fidelity, it is thought that the first genomes may have been smaller than today’s (62–64). On today’s tree of life, the CPR and DPANN clades both show small genome sizes, which, together with branch position, have been used to suggest that these organisms might be ancient - though recent analyses suggest a revision of these branch positions (3).

It is tempting to look at the scaling relationships presented here and consider where they might point to in terms of a minimal cell with a highly reduced genome, but [H] Coenzyme transport and metabolism, [C] Energy production and conversion, [I] Lipid transport and metabolism, [K] Transcription [M] Cell wall/membrane/envelope biogenesis, [T] Signal transduction mechanisms, and [U] Intracellular trafficking, secretion, and vesicular transport CPR and DPANN scaling relationships are unique, telling us that there are multiple ways to scale from this end of the genome size (Supplementary Table 2 and Supplementary Figure 5). On the other hand, the rest of the categories have the same scaling exponent, suggesting that these might be universal.

Most of the previous scaling analyses of bacterial physiology focus on the connection between cellular rates, processes, abundances, and cell size (9–11, 29). This is a natural connection, given that the scale of the system sets the values of various physical constraints. Such efforts have revealed how metabolic rates, growth rates, translational rates, and the abundances of all major macromolecules change with cell volume (10, 29). Genome size also changes with cell size systematically, which means that the relationships observed here also change with cell size, and this provides a way to compare how the abundance of functional categories changes with increasing size compared with how the overall physiology and macromolecular composition changes (10, 29).

Tracing back to the earliest cells, we are left with the question of how functional categories arrived in taxonomic clades: were they transferred horizontally from another group? or did they originate *de novo* within the group? In our analysis, functional groups associated with metabolic processes tend to have the highest variation in scaling exponents ([H] and [P] in figure 6). COG categories associated with metabolism also have been transferred the most between the domains (65), it may be that phyla-specific scaling patterns can be explained at least partially by the accretion of functions via horizontal gene transfer, though more detailed protein family-specific analyses are necessary to investigate repurposing/elaboration of existing genes or by de-novo birth and/or exaptation of other genes.

### Towards an understanding of evolution and scaling in biology

The relationships that we have described here in terms of the scaling of functions within genomes reveal fundamental regularities of genome organization. However, the underlying mechanisms behind these scaling relationships demand further explanation. Several simple mechanisms have been proposed for the scaling of certain categories. For example, the near quadratic scaling of regulatory genes suggests that every new gene addition to the genome comes with a pairwise consideration for regulation (the number of pairs grows like L^2^ given a fixed average gene length in a genome of total length L which is generally observed (12, 13, 22, 66). Nevertheless, such proposed mechanisms have not been elucidated for every COG category. In fact, even the scaling of macrophysiology - such as growth rates, metabolic rates, and macromolecular concentrations, with cell or genome size, does not give us a consistent way to explain the functional scaling relationships. For example, metabolic rate scales superlinearly with genome size in prokaryotes and sublinearly in eukaryotes (11). This overall scaling and shift between prokaryotes and eukaryotes is mirrored by the metabolic categories [C],[E], [G], and [H]. However, ribosome and accompanying translational macromolecule abundances scale superlinearly with genome size, but COG category [J] scales very sublinearly where the exponent is closer to zero. This could be because the ribosomal machinery is so conserved that there is not much expansion in the diversity of similar genes within [J].

Variability in scaling between phyla has been previously attributed to a dependence of genome size on the ease of horizontal gene transfer or genome plasticity (25); our observation of variability of scaling factors between phyla may suggest that genome plasticity is broadly different between these groups. Although we can offer explanations related to the regulation scaling, we cannot yet explain the full variability of exponents. Genome size correlates with cell size and cell size is a predictor of macro-physiology, so there is a hidden set of features that matter more than taxonomy.

While we include facets of scaling in biology not previously discussed, there are several things that deserve future attention, for example, copy number, for a more comprehensive understanding of evolution. Additionally, in our study, we focused exclusively on microbial organisms; future work which includes multicellular organisms will be valuable to link functional scaling with cell and organism plan. The differences in scaling observed using different projections of gene function also needs to be explained: Previous work has shown that EC categorization reveals universal scaling across the tree of life (17) compared with the many shifts in scaling observed here for COG categories. This points to deep questions about convergent evolution at the level of broad enzyme functionality (17) that is not observed at the level of physiological functionality. As the number of sequenced genomes and corresponding annotations increase over the years, we would expect our understanding of functional scaling to deepen and help us ask more relevant questions about organism design and functional evolution.

## AVAILABILITY

Jupyter notebooks are available at https://github.com/riddhi7/functional-scaling-across-genomes

## SUPPLEMENTARY DATA

Supplementary Data are available at NAR online.

## Acknowledgements

ACKNOWLEDGEMENTFUNDING

RG is funded by the ELSI Integrated Masters-PhD program. S.E.M. acknowledges support by NSF (Award No. 1724300) “Collaborative Research: Biochemical, Genetic, Metabolic, and Isotopic Constraints on an Ancient Thiobiosphere”

## CONFLICT OF INTEREST

The authors report no conflict of interest.

## Notes

### Competing Interest Statement

The authors have declared no competing interest.

