## Supplementary Figures for "Scaling of Protein Function Across the Tree of Life"

**[1] Earth-Life Science Institute, Tokyo Institute of Technology, Ookayama, Meguro-ku, 152-8550 Tokyo, Japan**

**[2] School of Life Sciences and Technology, Tokyo Institute of Technology, Tokyo, Ookayama, Meguro-ku, 152-8550 Tokyo, Japan**

**[3] The Santa Fe Institute, Santa Fe, NM, USA**

**[4] Blue Marble Space Institute of Science, Seattle, WA 98154, USA**

**[5] Center for Sustainable Resource Science, RIKEN, 2-1 Hirosawa, Wako, 351-0198 Saitama, Japan**

**This document contains supplementary figures:**

**Supplementary Figure 1**: Scaling in random orthogroups. Shuffled orthologs for a null test of the scaling exponents.
**Supplementary Figure 2:** Unibinned and binned power law fits.
**Supplementary Figure 3:** Phylogenetic distance plots.
**Supplementary Figure 4:** Phyla arranged in the increasing order of their exponents with 95% confidence intervals.
**Supplementary Figure 5:** Scaling comparison between CPR, DPANN, Asgard archaea, and Eukaryotes.

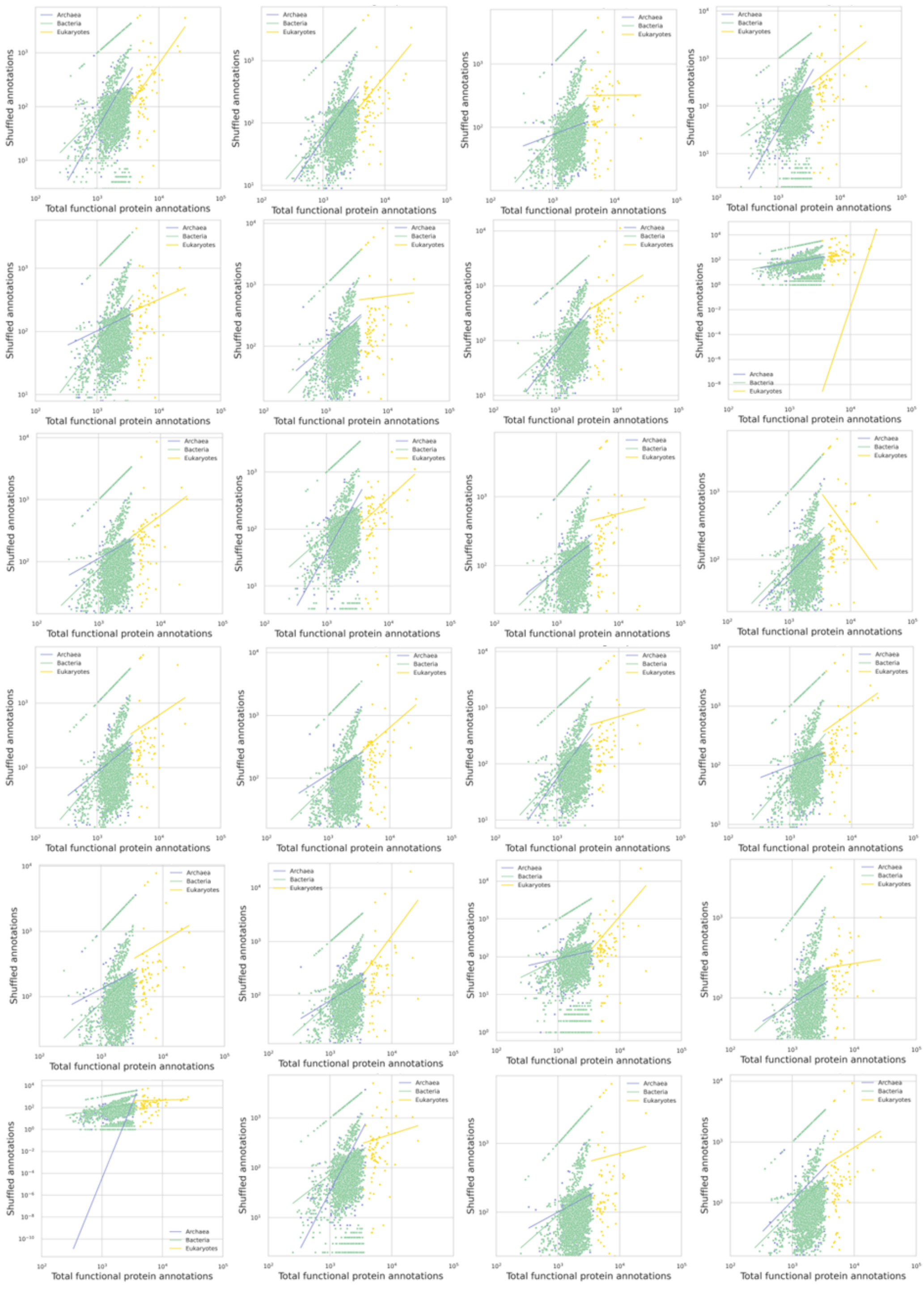


**Supplementary Figure 1: Scaling in random orthogroups. Shuffled orthologs for a null test of the scaling exponents.**

No specific patterns were observed when the orthogroups were shuffled.

### *1] Information Storage and Processing: COG categories A, B, J, K, and L*

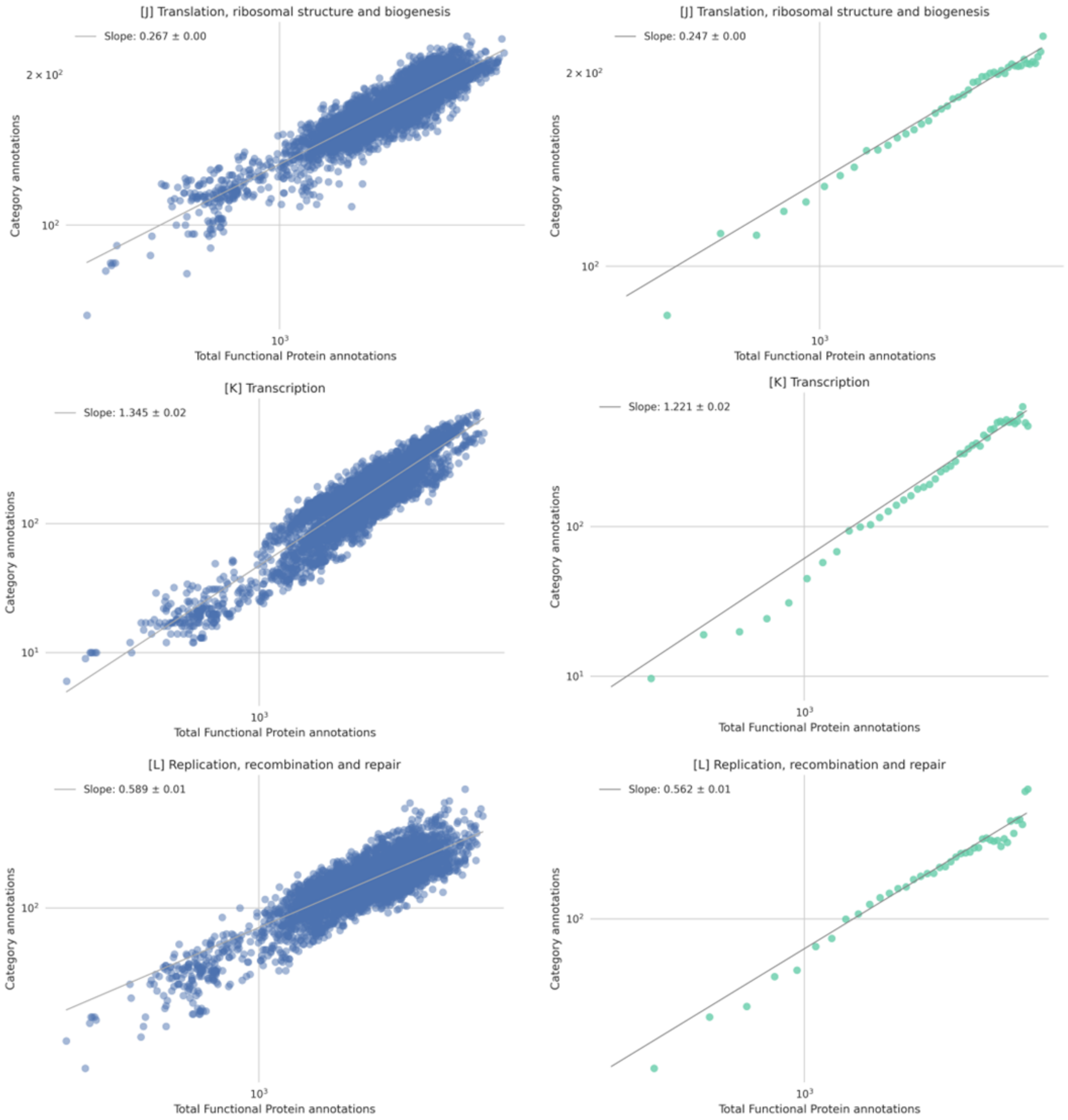

### 2] Cellular Processes and Signaling: COG categories D, M, N, O, T, U, V, W, Y, and Z

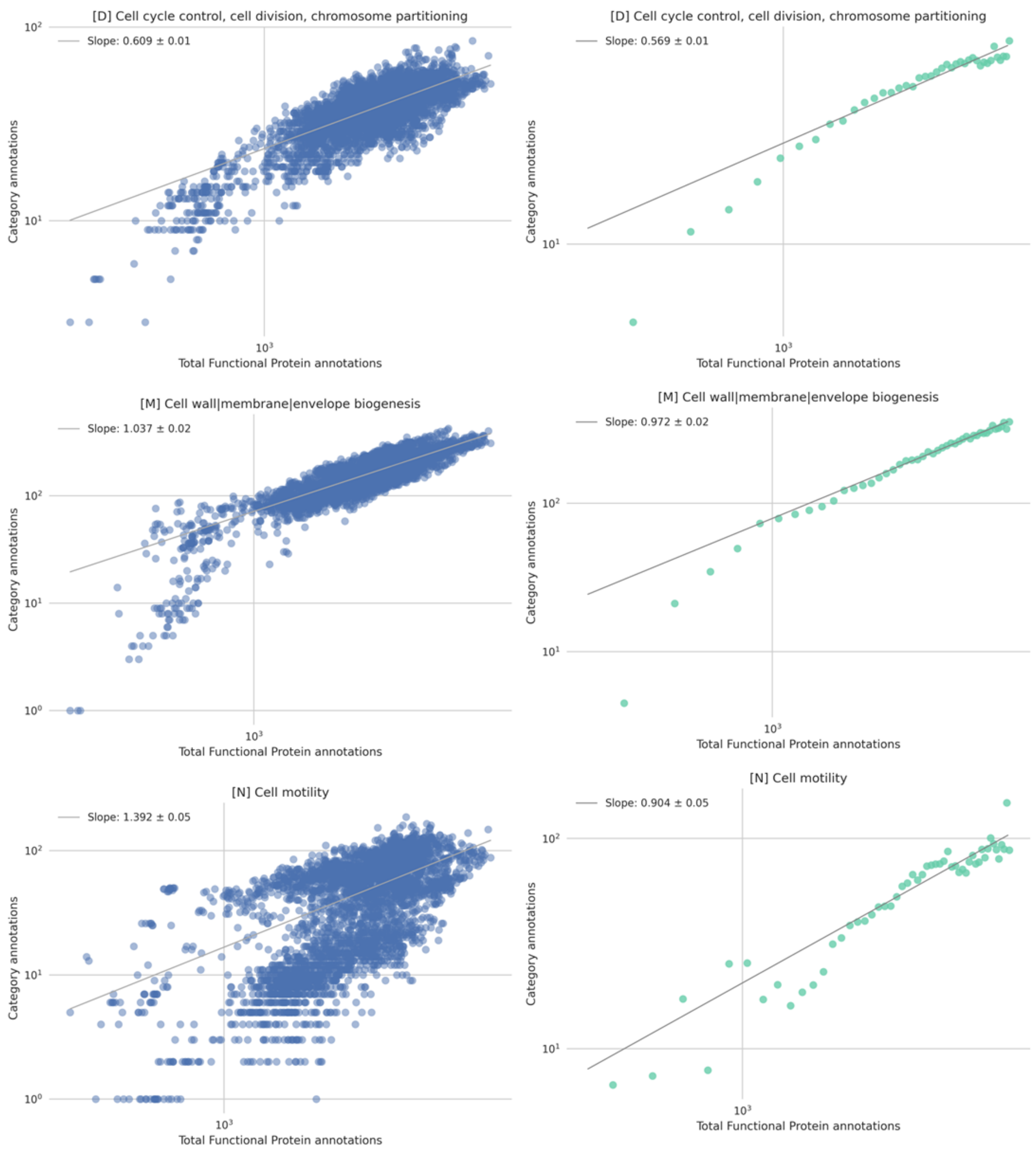

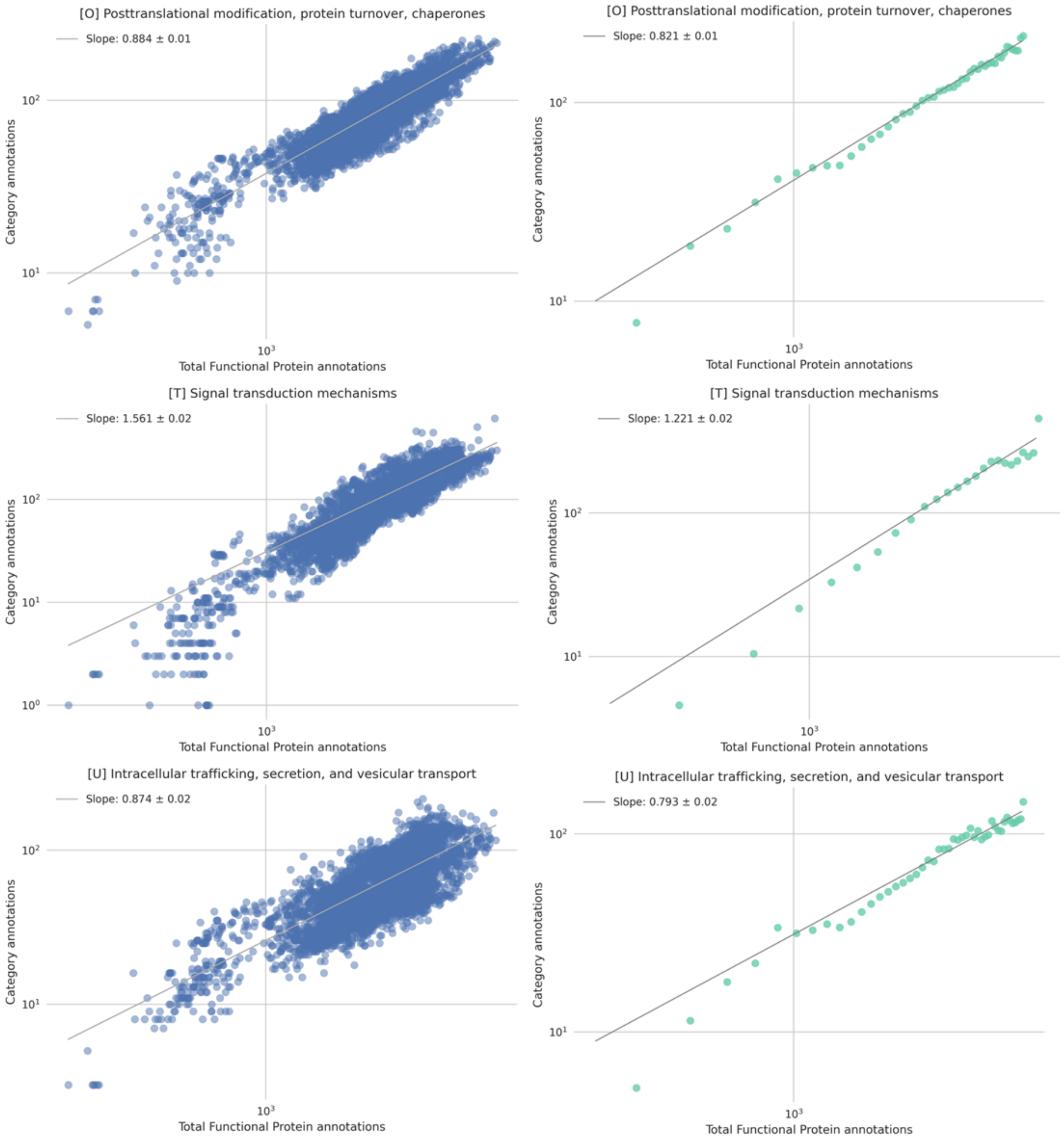
[O] Posttranslational modification, protein turnover, chaperones
Slope: 0.884 ± 0.01
Category annotations
Total Functional Protein annotations
[O] Posttranslational modification, protein turnover, chaperones
Slope: 0.821 ± 0.01
Category annotations
Total Functional Protein annotations
[T] Signal transduction mechanisms
Slope: 1.561 ± 0.02
Category annotations
Total Functional Protein annotations
[T] Signal transduction mechanisms
Slope: 1.221 ± 0.02
Category annotations
Total Functional Protein annotations
[U] Intracellular trafficking, secretion, and vesicular transport
Slope: 0.874 ± 0.02
Category annotations
Total Functional Protein annotations
[U] Intracellular trafficking, secretion, and vesicular transport
Slope: 0.793 ± 0.02
Category annotations
Total Functional Protein annotations

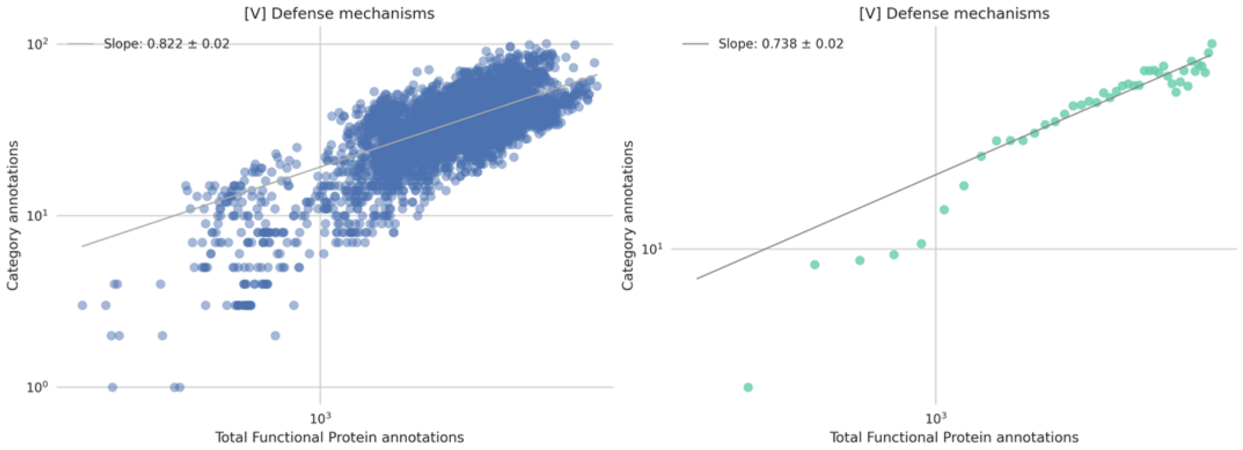

[V] Defense mechanisms
Slope: 0.822 ± 0.02
Category annotations
Total Functional Protein annotations
[V] Defense mechanisms
Slope: 0.738 ± 0.02
Category annotations
Total Functional Protein annotations

### 3] Metabolism: COG Categories C, E, F, G, H, I, P, and Q

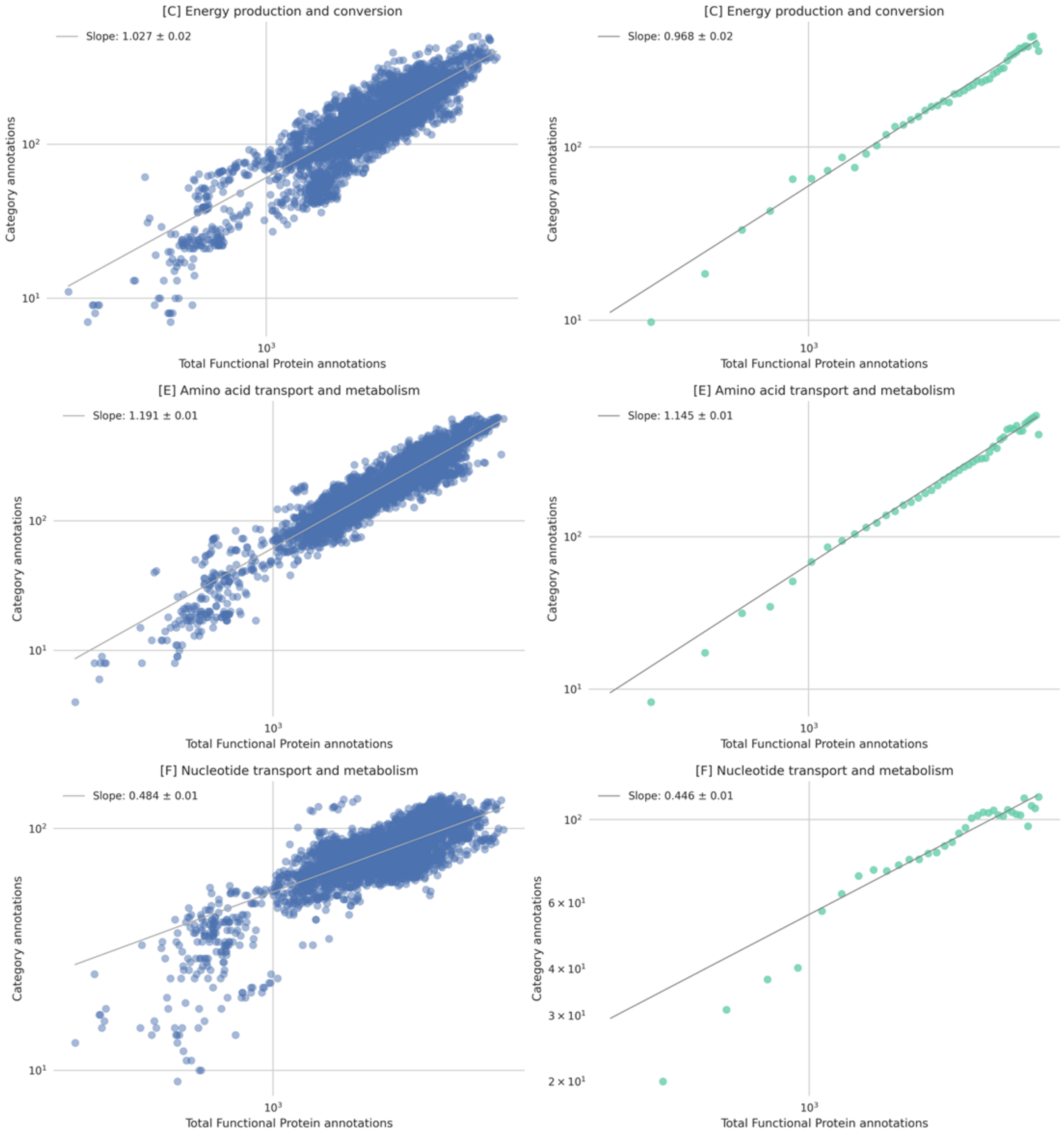

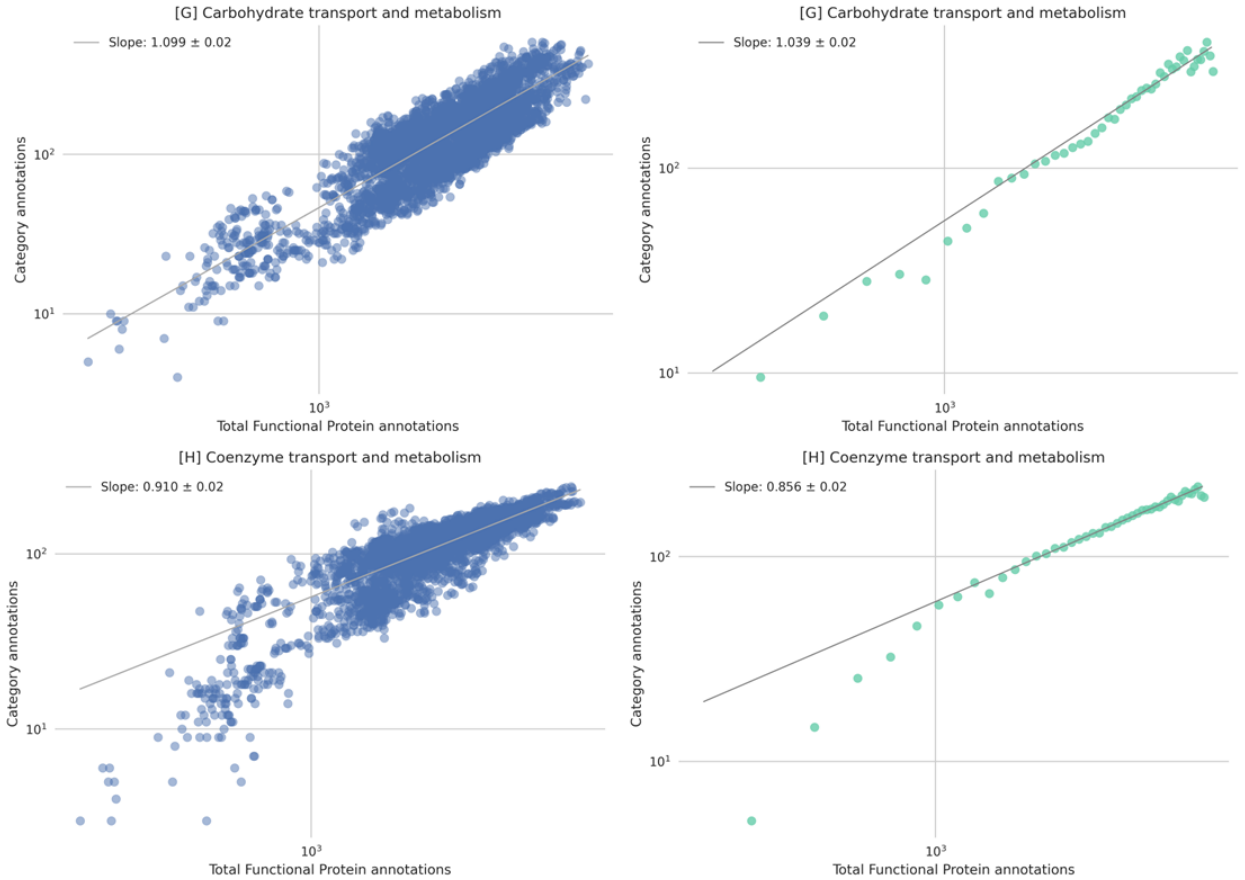
[G] Carbohydrate transport and metabolism
Slope: 1.099 ± 0.02
Category annotations
Total Functional Protein annotations
[G] Carbohydrate transport and metabolism
Slope: 1.039 ± 0.02
Category annotations
Total Functional Protein annotations
[H] Coenzyme transport and metabolism
Slope: 0.910 ± 0.02
Category annotations
Total Functional Protein annotations
[H] Coenzyme transport and metabolism
Slope: 0.856 ± 0.02
Category annotations
Total Functional Protein annotations

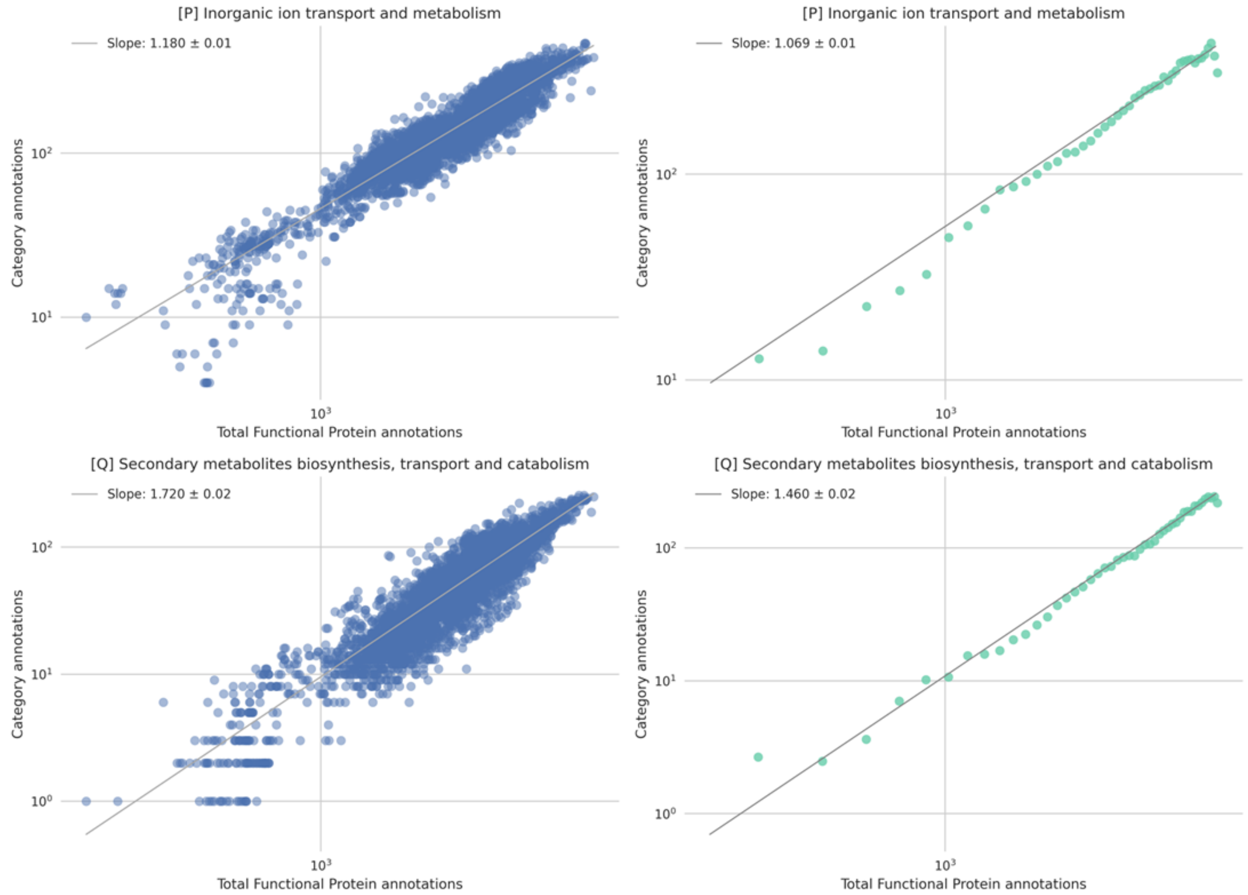


**Supplementary Figure 2: Unbinned and binned power law fits.** Axes are in log scale.

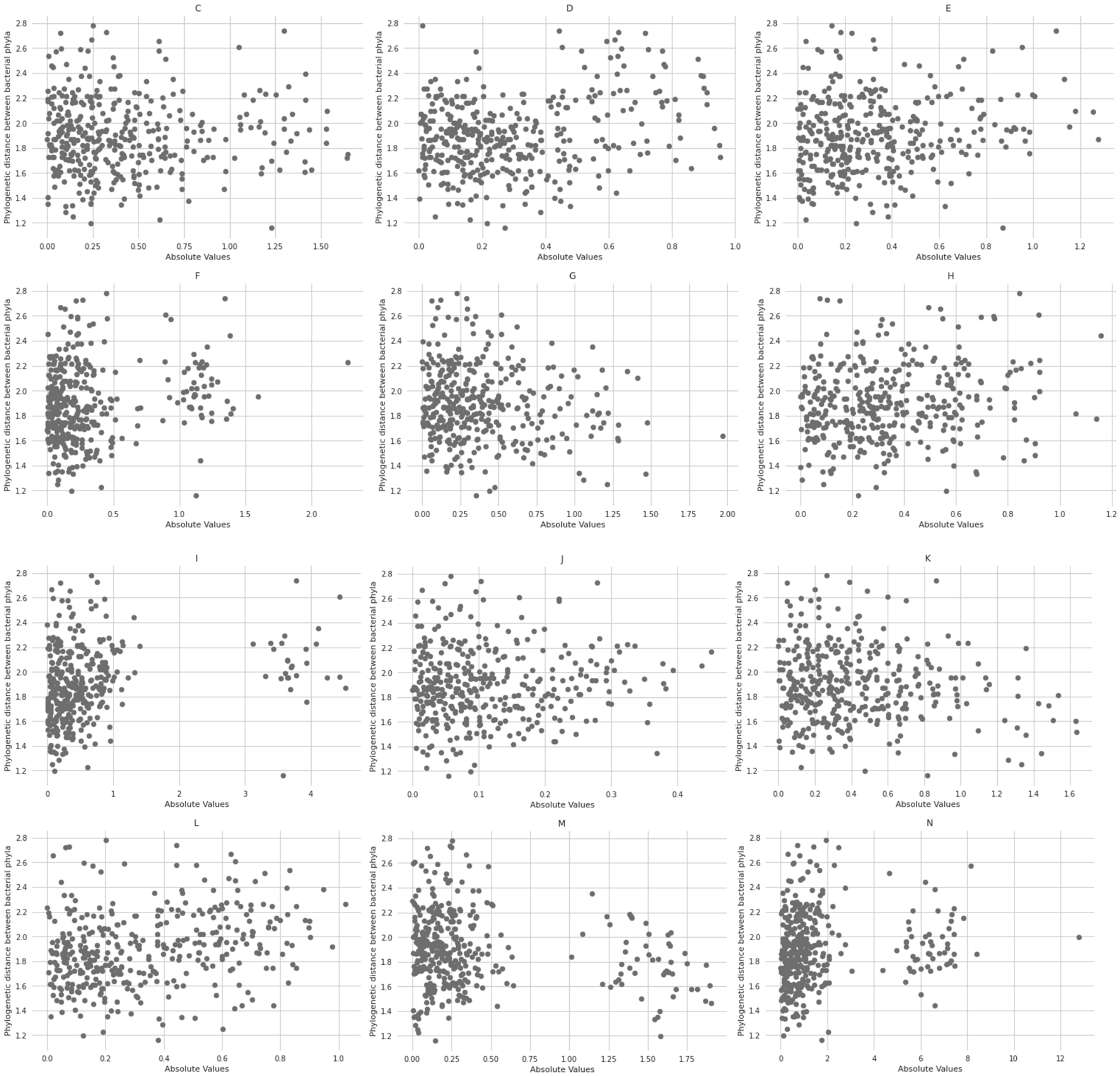

C
D
E
F
G
H
I
J
K
L
M
N
Phylogenetic distance between bacterial phyla
Absolute Values

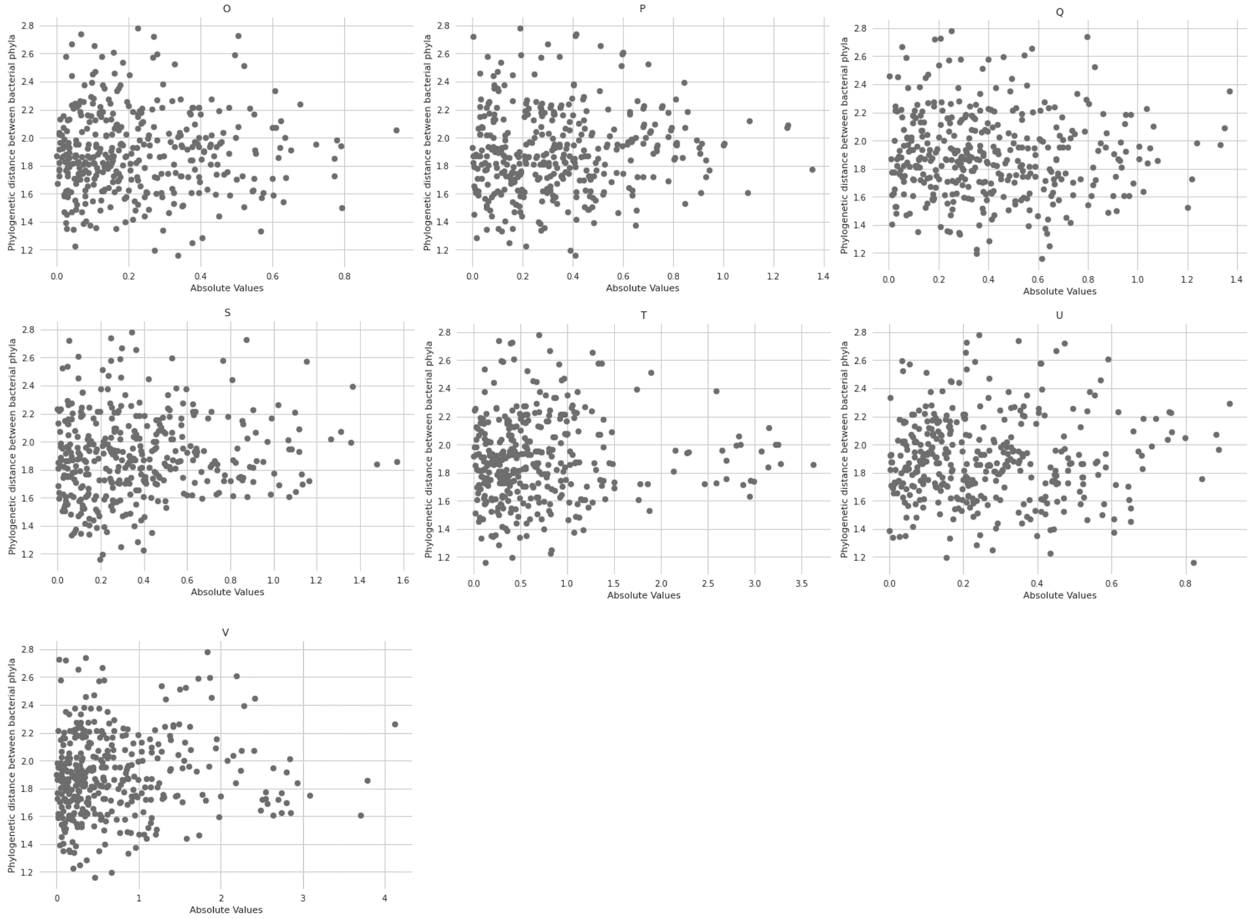


**Supplementary Figure 3: Phylogenetic distance plots.** Absolute value is the difference between the exponents of the respective phyla.

### 1] Information Storage and Processing: COG categories A, B, J, K, and L

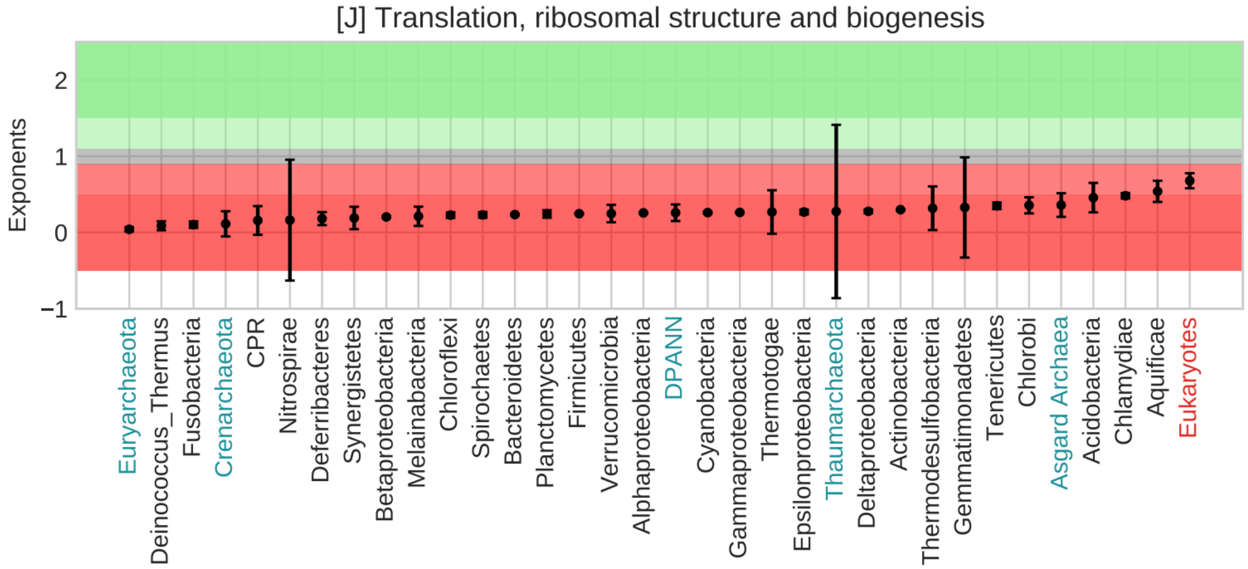


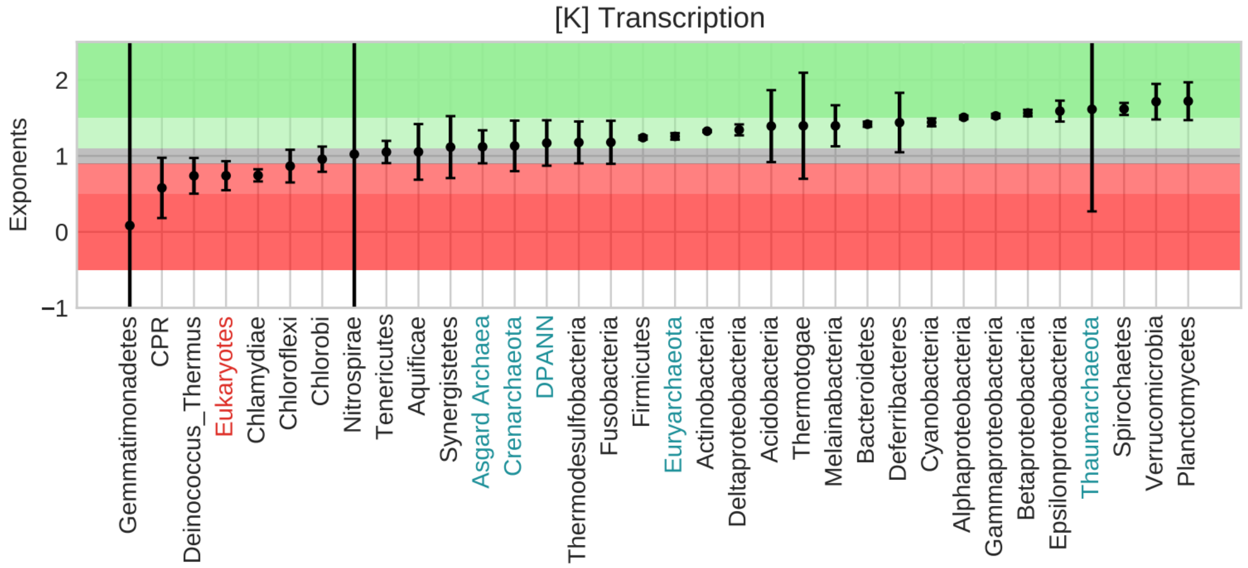

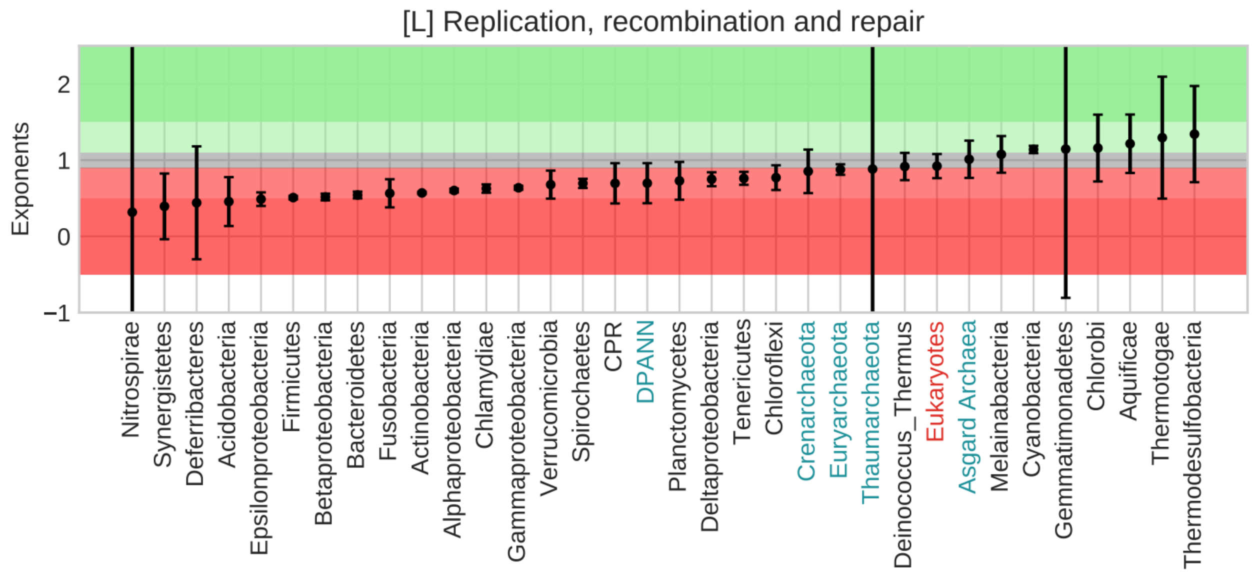


### 2] Cellular Processes and Signaling: COG categories D, M, N, O, T, U, V, W, Y, and Z

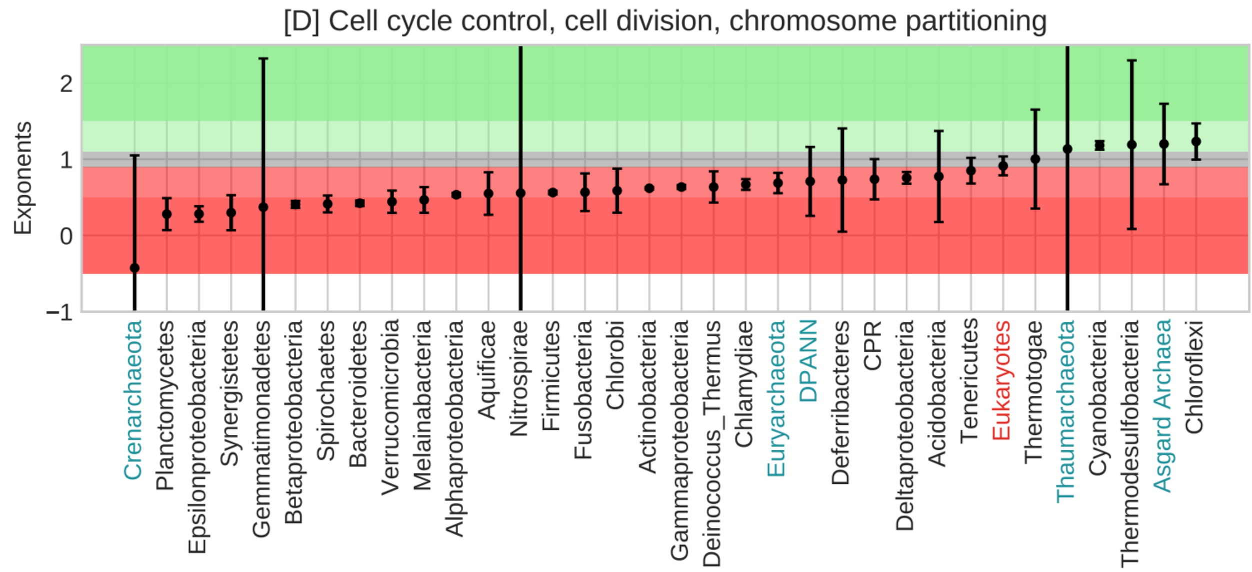

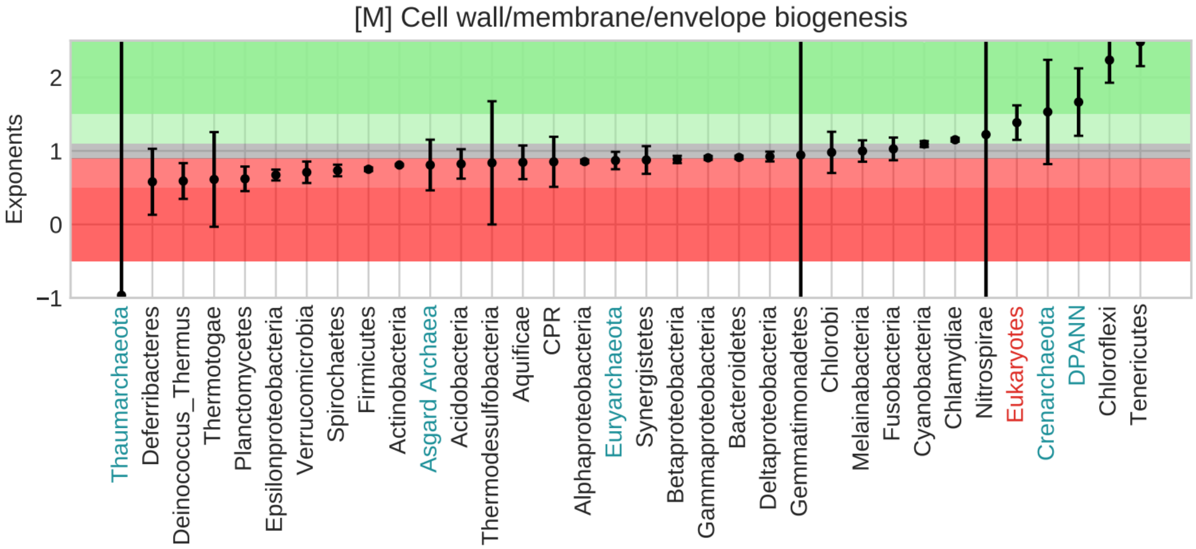
[M] Cell wall/membrane/envelope biogenesis
Exponents
2
1
0
−1
Thaumarchaeota
Deferribacteres
Deinococcus_Thermus
Thermotogae
Planctomycetes
Epsilonproteobacteria
Verrucomicrobia
Spirochaetes
Firmicutes
Actinobacteria
Asgard Archaea
Acidobacteria
Thermodesulfobacteria
Aquificae
CPR
Alphaproteobacteria
Euryarchaeota
Synergistetes
Betaproteobacteria
Gammaproteobacteria
Bacteroidetes
Deltaproteobacteria
Gemmatimonadetes
Chlorobi
Melainabacteria
Fusobacteria
Cyanobacteria
Chlamydiae
Nitrospirae
Eukaryotes
Crenarchaeota
DPANN
Chloroflexi
Tenericutes

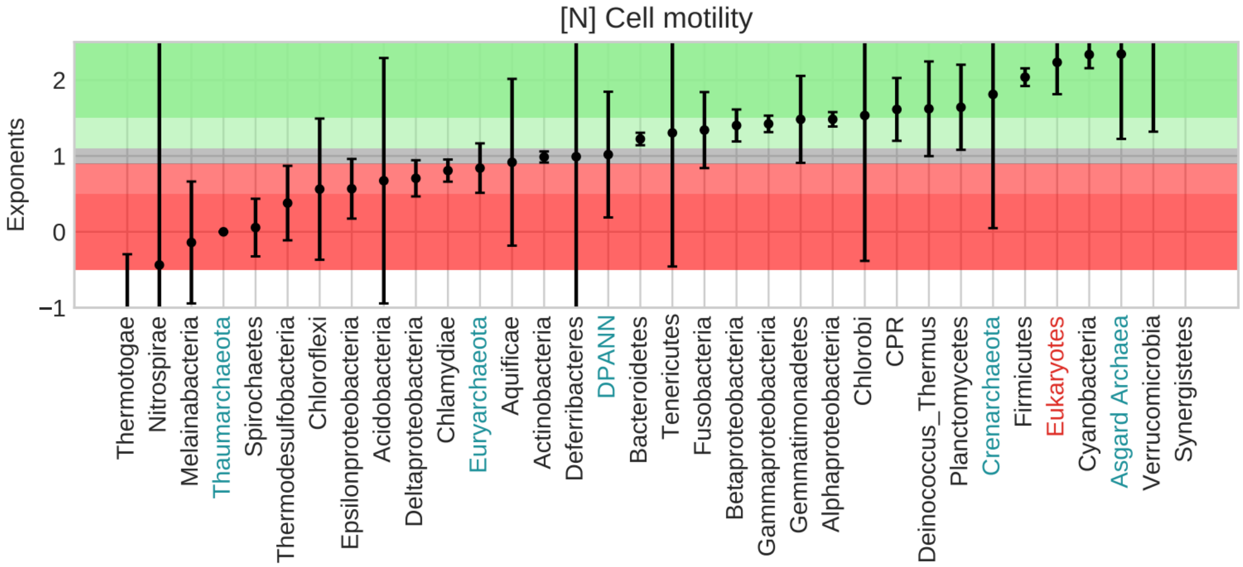
[N] Cell motility
Exponents
2
1
0
−1
Thermotogae
Nitrospirae
Melainabacteria
Thaumarchaeota
Spirochaetes
Thermodesulfobacteria
Chloroflexi
Epsilonproteobacteria
Acidobacteria
Deltaproteobacteria
Chlamydiae
Euryarchaeota
Aquificae
Actinobacteria
Deferribacteres
DPANN
Bacteroidetes
Tenericutes
Fusobacteria
Betaproteobacteria
Gammaproteobacteria
Gemmatimonadetes
Alphaproteobacteria
Chlorobi
CPR
Deinococcus_Thermus
Planctomycetes
Crenarchaeota
Firmicutes
Eukaryotes
Cyanobacteria
Asgard Archaea
Verrucomicrobia
Synergistetes

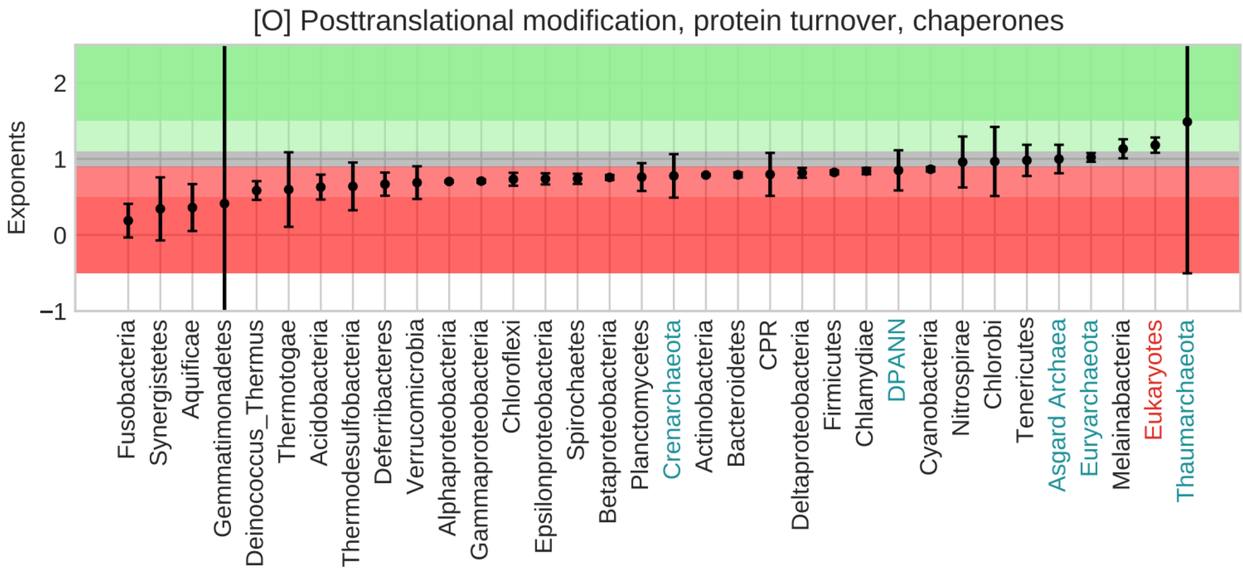

[O] Posttranslational modification, protein turnover, chaperones
Exponents
2
1
0
−1
Fusobacteria
Synergistetes
Aquificae
Gemmatimonadetes
Deinococcus_Thermus
Thermotogae
Acidobacteria
Thermodesulfobacteria
Deferribacteres
Verrucomicrobia
Alphaproteobacteria
Gammaproteobacteria
Chloroflexi
Epsilonproteobacteria
Spirochaetes
Betaproteobacteria
Planctomycetes
Crenarchaeota
Actinobacteria
Bacteroidetes
CPR
Deltaproteobacteria
Firmicutes
Chlamydiae
DPANN
Cyanobacteria
Nitrospirae
Chlorobi
Tenericutes
Asgard Archaea
Euryarchaeota
Melainabacteria
Eukaryotes
Thaumarchaeota


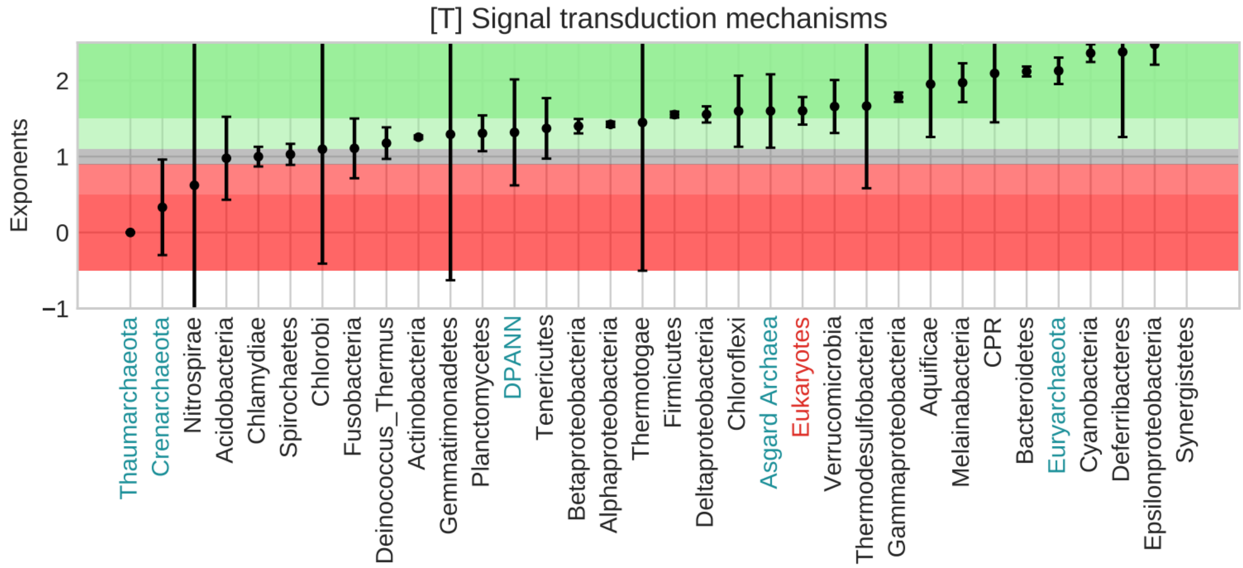

[T] Signal transduction mechanisms
Exponents
2
1
0
−1
Thaumarchaeota
Crenarchaeota
Nitrospirae
Acidobacteria
Chlamydiae
Spirochaetes
Chlorobi
Fusobacteria
Deinococcus_Thermus
Actinobacteria
Gemmatimonadetes
Planctomycetes
DPANN
Tenericutes
Betaproteobacteria
Alphaproteobacteria
Thermotogae
Firmicutes
Deltaproteobacteria
Chloroflexi
Asgard Archaea
Eukaryotes
Verrucomicrobia
Thermodesulfobacteria
Gammaproteobacteria
Aquificae
Melainabacteria
CPR
Bacteroidetes
Euryarchaeota
Cyanobacteria
Deferribacteres
Epsilonproteobacteria
Synergistetes

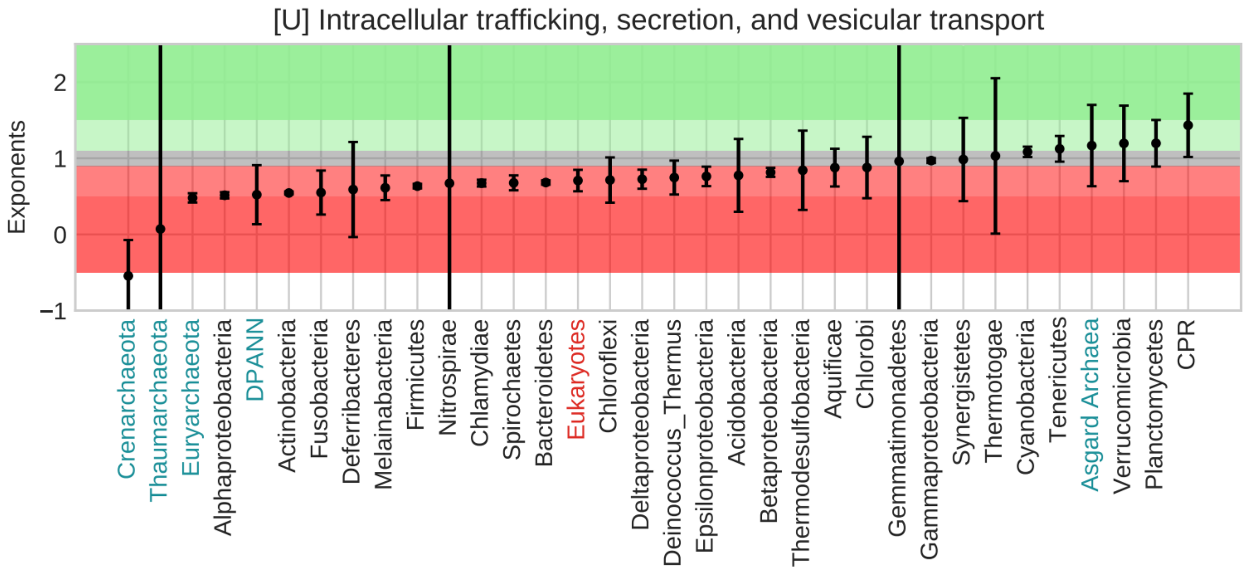
[U] Intracellular trafficking, secretion, and vesicular transport
Exponents
2
1
0
−1
Crenarchaeota
Thaumarchaeota
Euryarchaeota
Alphaproteobacteria
DPANN
Actinobacteria
Fusobacteria
Deferribacteres
Melainabacteria
Firmicutes
Nitrospirae
Chlamydiae
Spirochaetes
Bacteroidetes
Eukaryotes
Chloroflexi
Deltaproteobacteria
Deinococcus_Thermus
Epsilonproteobacteria
Acidobacteria
Betaproteobacteria
Thermodesulfobacteria
Aquificae
Chlorobi
Gemmatimonadetes
Gammaproteobacteria
Synergistetes
Thermotogae
Cyanobacteria
Tenericutes
Asgard Archaea
Verrucomicrobia
Planctomycetes
CPR

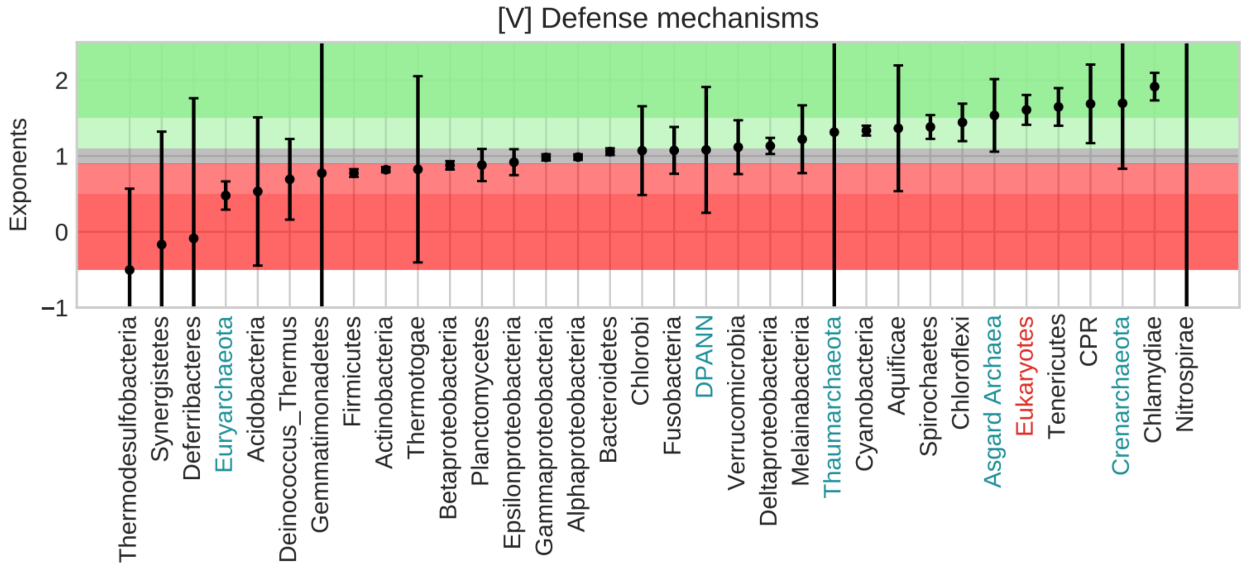
[V] Defense mechanisms
Exponents
2
1
0
−1
Thermodesulfobacteria
Synergistetes
Deferribacteres
Euryarchaeota
Acidobacteria
Deinococcus_Thermus
Gemmatimonadetes
Firmicutes
Actinobacteria
Thermotogae
Betaproteobacteria
Planctomycetes
Epsilonproteobacteria
Gammaproteobacteria
Alphaproteobacteria
Bacteroidetes
Chlorobi
Fusobacteria
DPANN
Verrucomicrobia
Deltaproteobacteria
Melainabacteria
Thaumarchaeota
Cyanobacteria
Aquificae
Spirochaetes
Chloroflexi
Asgard Archaea
Eukaryotes
Tenericutes
CPR
Crenarchaeota
Chlamydiae
Nitrospirae

### *3] Metabolism: COG Categories C, E, F, G, H, I, P, and Q*

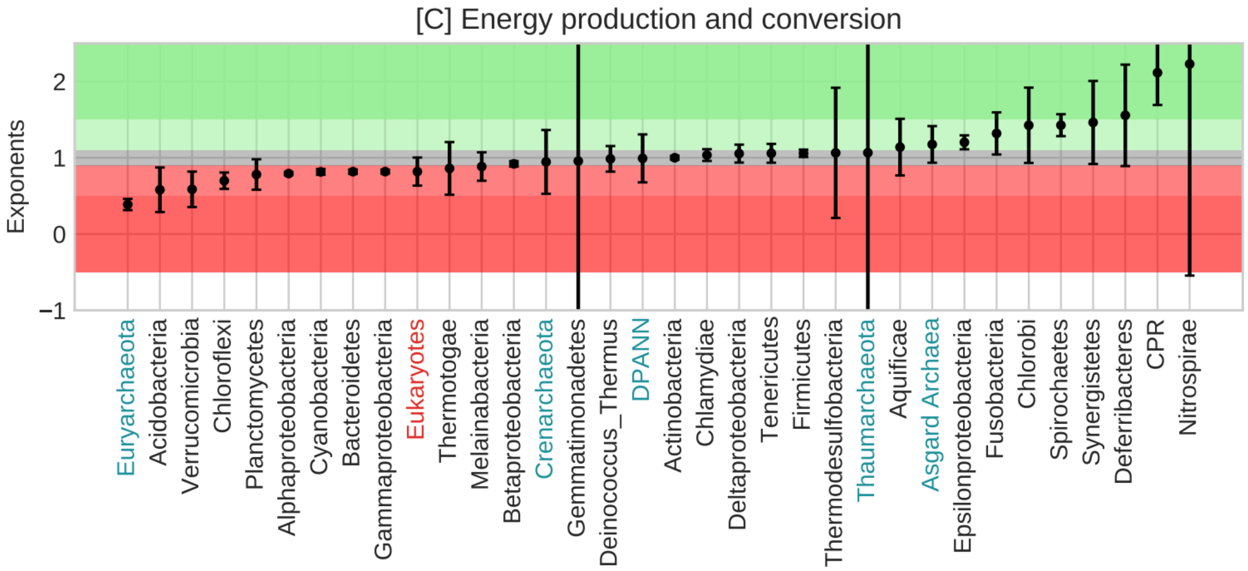


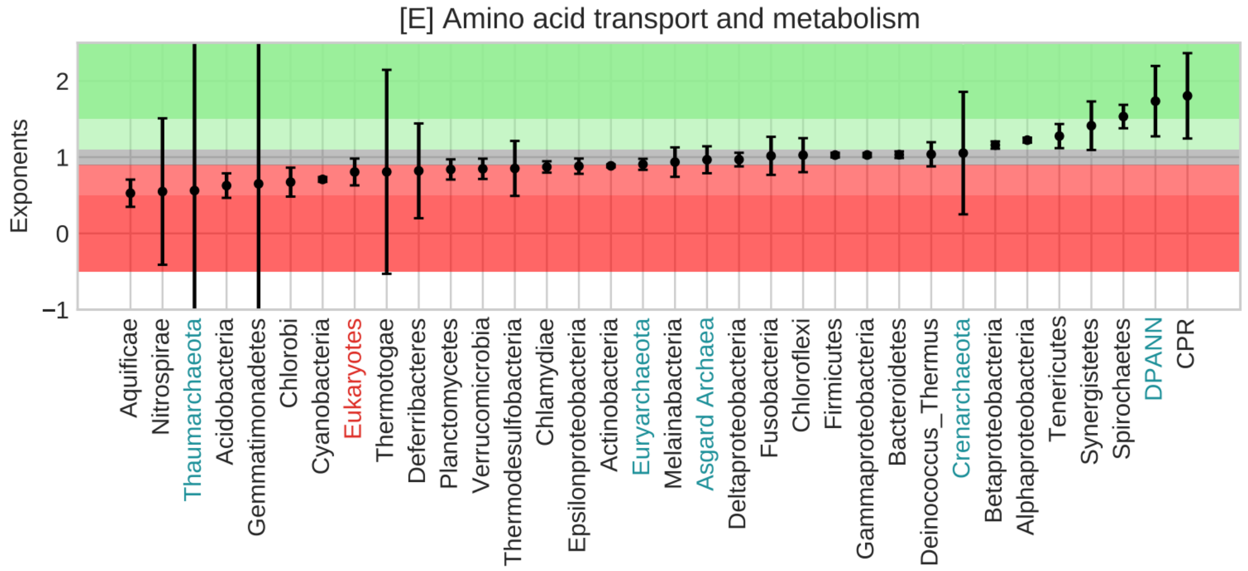

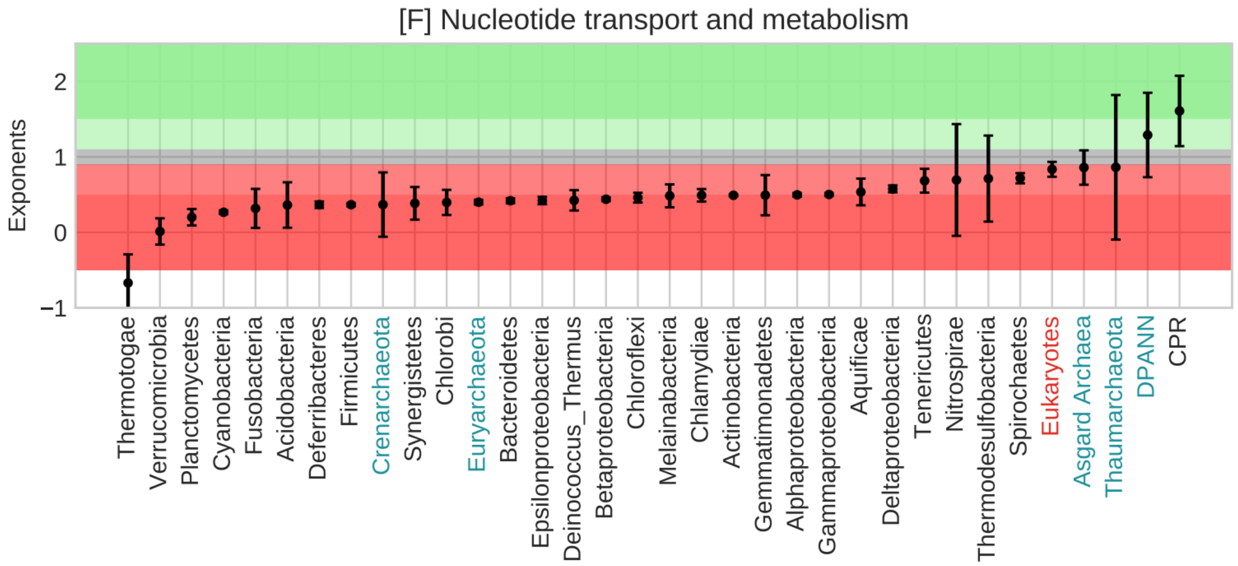
[F] Nucleotide transport and metabolism
Exponents
2
1
0
−1
Thermotogae
Verrucomicrobia
Planctomycetes
Cyanobacteria
Fusobacteria
Acidobacteria
Deferribacteres
Firmicutes
Crenarchaeota
Synergistetes
Chlorobi
Euryarchaeota
Bacteroidetes
Epsilonproteobacteria
Deinococcus_Thermus
Betaproteobacteria
Chloroflexi
Melainabacteria
Chlamydiae
Actinobacteria
Gemmatimonadetes
Alphaproteobacteria
Gammaproteobacteria
Aquificae
Deltaproteobacteria
Tenericutes
Nitrospirae
Thermodesulfobacteria
Spirochaetes
Eukaryotes
Asgard Archaea
Thaumarchaeota
DPANN
CPR

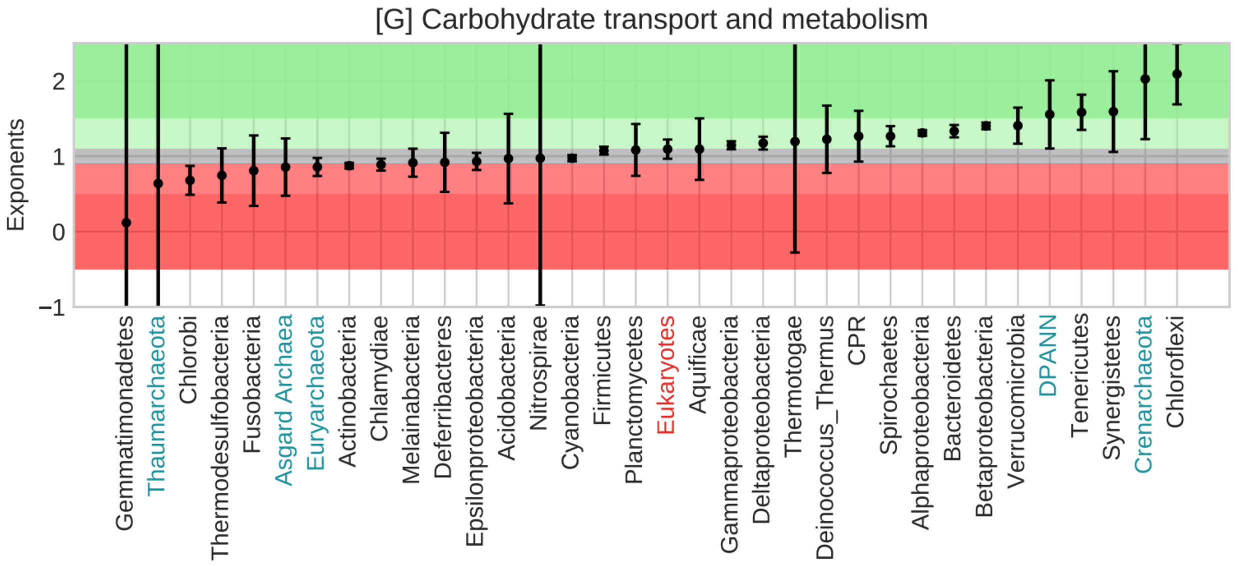
[G] Carbohydrate transport and metabolism
Exponents
2
1
0
−1
Gemmatimonadetes
Thaumarchaeota
Chlorobi
Thermodesulfobacteria
Fusobacteria
Asgard Archaea
Euryarchaeota
Actinobacteria
Chlamydiae
Melainabacteria
Deferribacteres
Epsilonproteobacteria
Acidobacteria
Nitrospirae
Cyanobacteria
Firmicutes
Planctomycetes
Eukaryotes
Aquificae
Gammaproteobacteria
Deltaproteobacteria
Thermotogae
Deinococcus_Thermus
CPR
Spirochaetes
Alphaproteobacteria
Bacteroidetes
Betaproteobacteria
Verrucomicrobia
DPANN
Tenericutes
Synergistetes
Crenarchaeota
Chloroflexi

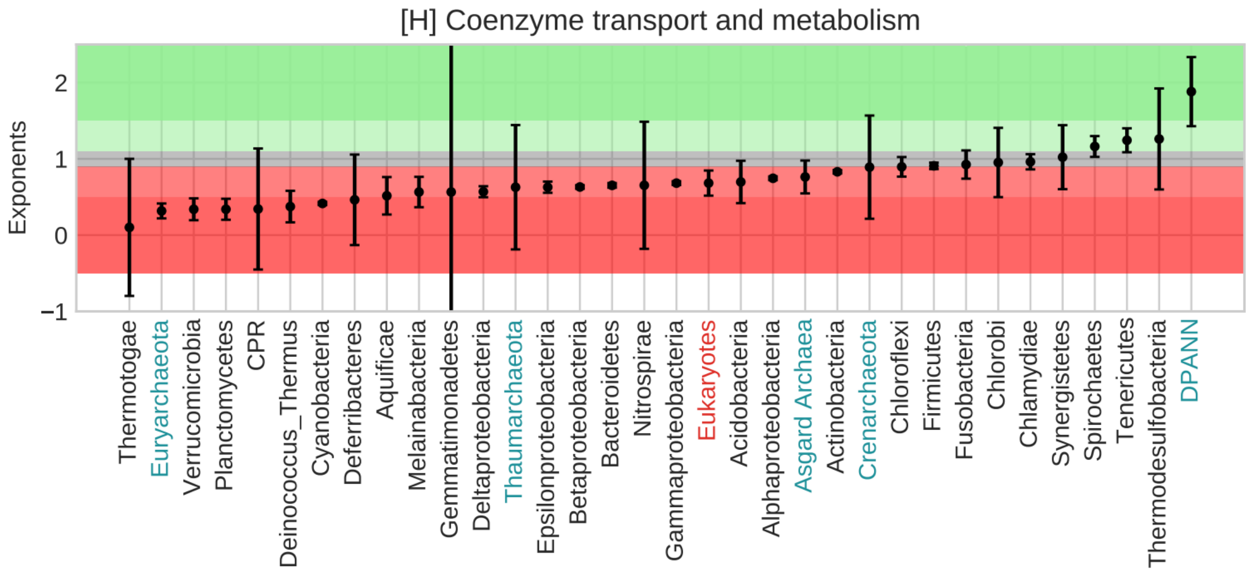
[H] Coenzyme transport and metabolism
Exponents
2
1
0
−1
Thermotogae
Euryarchaeota
Verrucomicrobia
Planctomycetes
CPR
Deinococcus_Thermus
Cyanobacteria
Deferribacteres
Aquificae
Melainabacteria
Gemmatimonadetes
Deltaproteobacteria
Thaumarchaeota
Epsilonproteobacteria
Betaproteobacteria
Bacteroidetes
Nitrospirae
Gammaproteobacteria
Eukaryotes
Acidobacteria
Alphaproteobacteria
Asgard Archaea
Actinobacteria
Crenarchaeota
Chloroflexi
Firmicutes
Fusobacteria
Chlorobi
Chlamydiae
Synergistetes
Spirochaetes
Tenericutes
Thermodesulfobacteria
DPANN

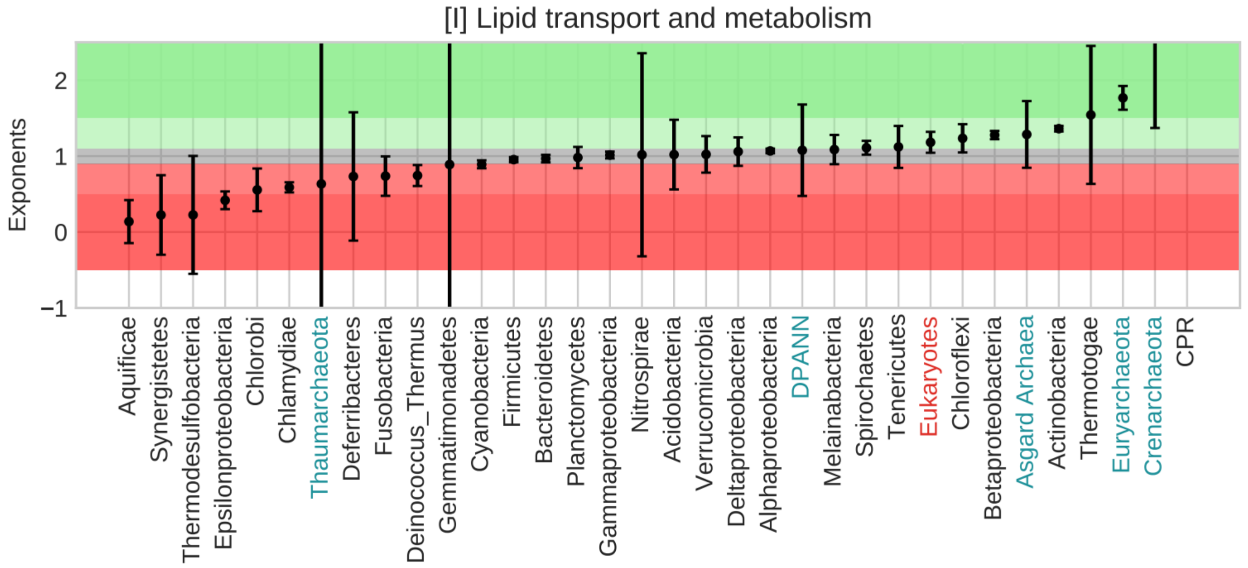
[I] Lipid transport and metabolism
Exponents
2
1
0
−1
Aquificae
Synergistetes
Thermodesulfobacteria
Epsilonproteobacteria
Chlorobi
Chlamydiae
Thaumarchaeota
Deferribacteres
Fusobacteria
Deinococcus_Thermus
Gemmatimonadetes
Cyanobacteria
Firmicutes
Bacteroidetes
Planctomycetes
Gammaproteobacteria
Nitrospirae
Acidobacteria
Verrucomicrobia
Deltaproteobacteria
Alphaproteobacteria
DPANN
Melainabacteria
Spirochaetes
Tenericutes
Eukaryotes
Chloroflexi
Betaproteobacteria
Asgard Archaea
Actinobacteria
Thermogae
Euryarchaeota
Crenarchaeota
CPR

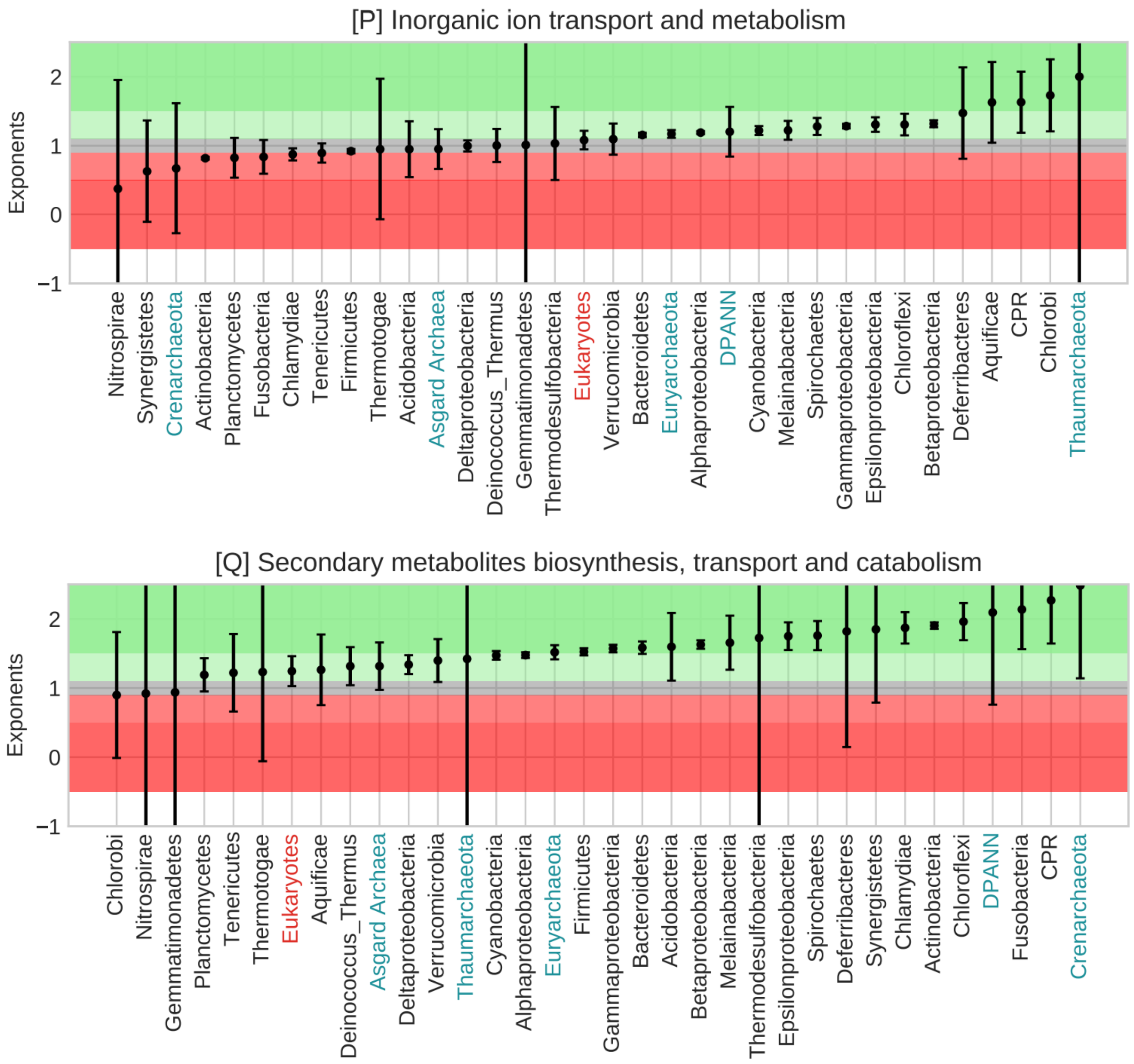


**Supplementary Figure 4:** Phyla arranged in the increasing order of their exponents with 95% confidence intervals. Background horizontal span colors signify scaling. From dark green to dark red, meaning superlinear scaling to sublinear scaling. The grey span in between signifies linear scaling.
X-axis labels: Bacterial phyla in black, archaeal phyla in cyan, and Eukaryotes in red.

### *1] Information Storage and Processing: COG categories A, B, J, K, and L*

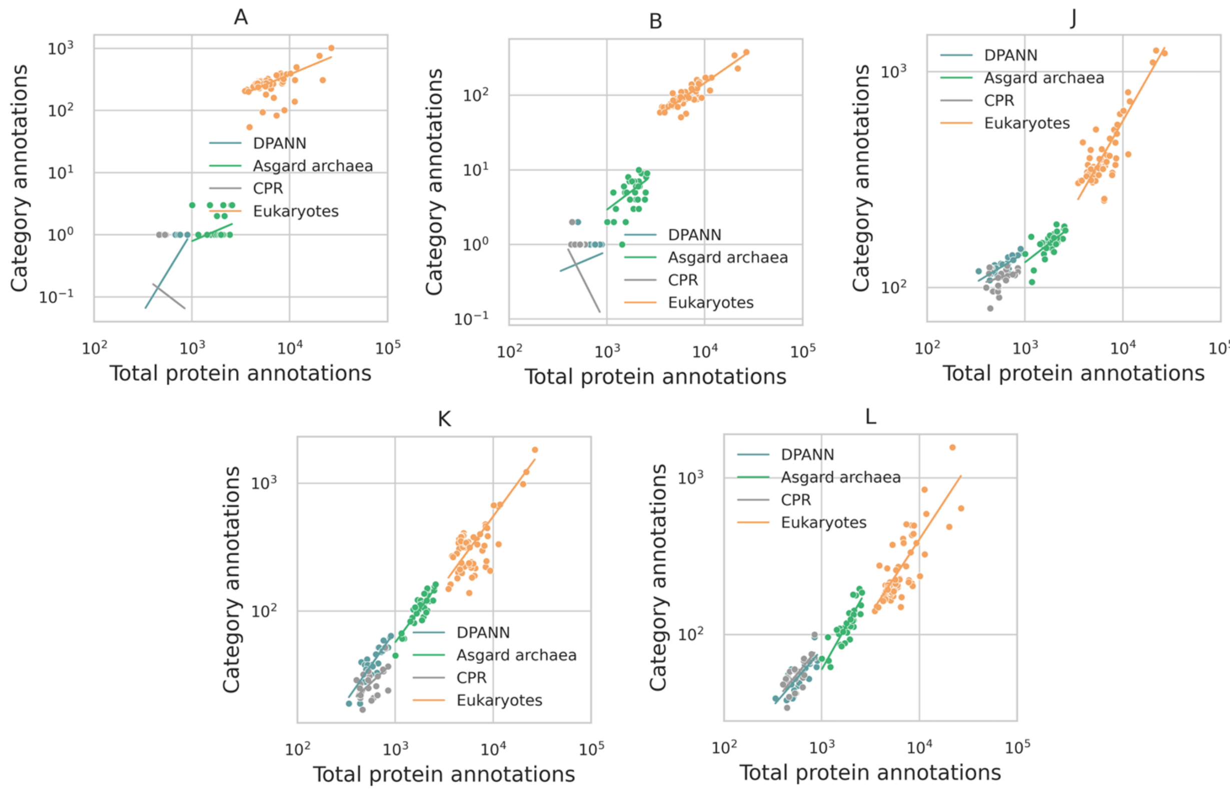


### *2] Cellular Processes and Signaling: COG categories D, M, N, O, T, U, V, W, Y, and Z*

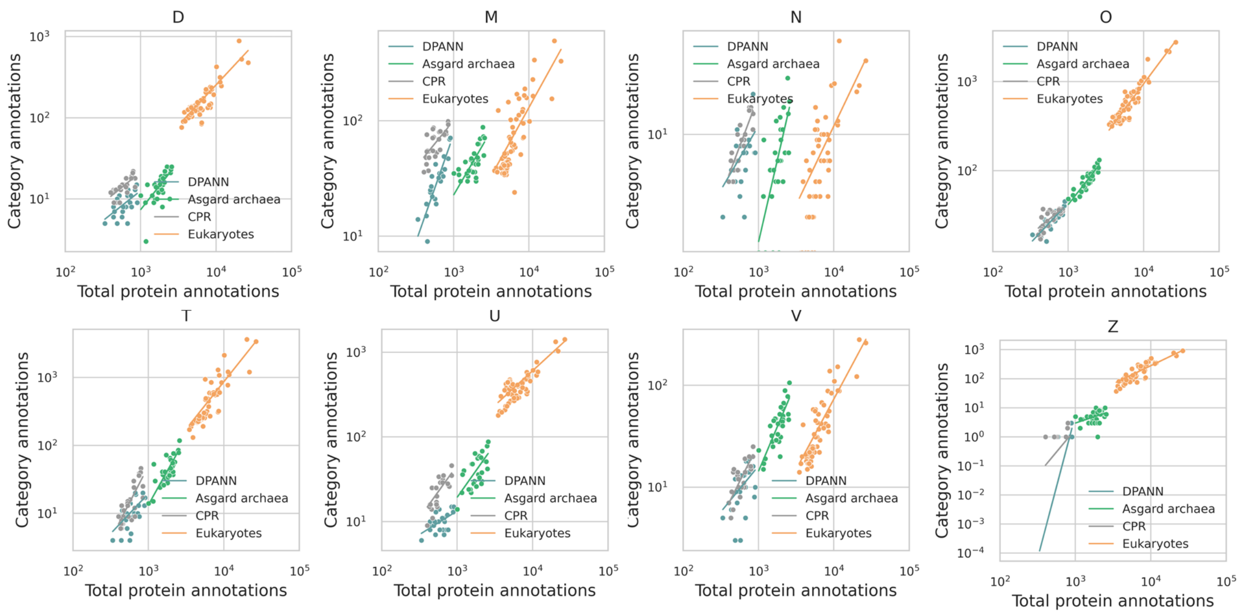

### *3] Metabolism: COG Categories C, E, F, G, H, I, P, and Q*

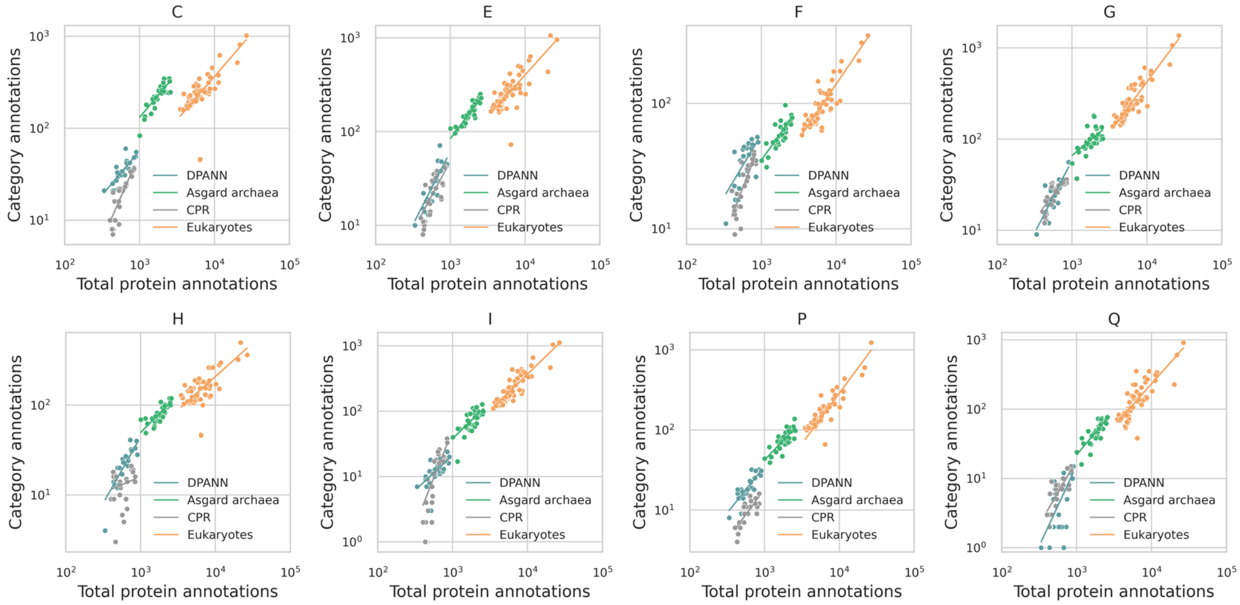


**Supplementary Figure 5: Scaling comparison between CPR, DPANN, Asgard archaea, and Eukaryotes**
